# Structural survey of HIF-2α reveals regulation of its subcellular localization and protein interactome

**DOI:** 10.64898/2026.05.04.722697

**Authors:** Tomas Gregor, Sinan Karakaya, Anja Zethraeus, Stina Östenberg, Elina Fredlund, Emma U. Hammarlund, Jenny Hansson, Jared Rosenblum, Sofie Mohlin

## Abstract

Hypoxia-inducible factor 2α (HIF-2α) is a central regulator of cellular homeostasis and a known oncogenic driver in multiple cancers. Although HIF-2α is canonically defined as a nuclear transcription factor, its cytoplasmic presence and non-canonical functions remain poorly understood. Here, we performed a structural survey of HIF-2α to determine the mechanisms underlying subcellular localization, protein abundance, and activity using a deletion-construct library, transcriptional assays, and *in vivo* xenograft models. We found that the oxygen-dependent degradation domain (ODD), the N-terminal intrinsically disordered region (n-IDR) and the N-terminal transactivation domain (NTAD) promote cytoplasmic localization, whereas the C-terminal IDR drives nuclear accumulation. Surprisingly, we found that HIF-2α nuclear localization occurs also in the absence of PAS A and B, the domains required for ARNT (HIF-1β) dimerization, resolving the long-standing question in the field. These data suggest a dominant role for non-canonical cytoplasmic mechanisms in HIF-2α-driven tumorigenesis. Strikingly, neither NTAD nor the C-terminal CTAD was required for tumor growth *in vivo*, in coherence with our transcriptional assays indicating that CTAD is dispensable for transactivation and NTAD functions as a suppressor rather than an activator of transcription. Proteomic analyses reveal HIF-2α interactions with regulators of mitochondrial function, translation initiation, RNA splicing, vesicular transport, and DNA replication. Together, these findings uncover previously unrecognized structural and functional complexity of HIF-2α compartmentalization and expand its role beyond canonical transcriptional regulation.

## Introduction

Adaptation to fluctuating oxygen levels is critical for maintaining cellular and tissue homeostasis during development and throughout adult human life (*1*). Central to this process are the hypoxia-inducible factors (HIFs), intracellular oxygen sensors that coordinate transcriptional programs in response to changes in oxygen availability. Initial studies of HIFs discovered their critical canonical role as nuclear transcription factors (*2*). However, additional studies have revealed that important aspects of their role in biology remain incompletely understood (*3–5*). Here, we uncover previously uncharacterized functions of the HIF family member HIF-2α (encoded by *EPAS1*), expanding the current view of HIF biology.

The HIF family comprises three α-isoforms, HIF-1α, HIF-2α, and HIF-3α, that exhibit distinct temporal and tissue-specific roles (*1*). HIF-1α is broadly expressed and primarily mediates acute hypoxic responses, with its levels declining rapidly once the acute phases settle. In contrast, HIF-2α activity is more tightly regulated and emerges after the initial HIF-1α response subsides, supporting sustained cellular adaptation. We previously demonstrated that precise control of HIF-2α expression is essential for trunk neural crest stem cell development during embryogenesis (*6*) and that high cytoplasmic HIF-2α predicts poor prognosis in sympathetic paraganglioma (*4*). Elevated HIF-2α expression is likewise associated with adverse outcomes across multiple malignancies, including neuroblastoma, pheochromocytoma, renal cell carcinoma, glioma, chondrosarcoma, melanoma, and cancers of the breast, head and neck, liver, prostate, lung, and pancreas (*7*). In neuroblastoma, high-intensity protein expression of HIF-2α, but not HIF-1α, in a distinct subpopulation of cells has been specifically associated with metastatic progression and poor overall survival (*8–10*).

Under physiological oxygen levels (physioxia), HIF-α proteins are continuously synthesized but rapidly targeted for degradation via von Hippel-Lindau (VHL)-mediated ubiquitination following oxygen-dependent prolyl hydroxylation by prolyl hydroxylase domain (PHD/EGLN) enzymes (*2*, *11*). HIF-α activity is further constrained by factor-inhibiting HIF-1 (FIH1), which catalyzes oxygen-dependent asparagine hydroxylation within the C-terminal transactivation domain, thereby preventing interaction with the transcriptional co-activators CBP/p300 (*12*, *13*). When oxygen availability drops below physioxic levels (hypoxia), hydroxylation of HIF-α cannot occur since molecular oxygen is an obligate co-substrate for both PHD and FIH enzymes. As a result, PHDs are unable to modify HIF-α, VHL cannot recognize or bind the protein, and HIF-α escapes proteasomal degradation. Typically, stabilized HIF-α then accumulates, translocates to the nucleus, and activates transcription of hypoxia response element (HRE)-containing genes through heterodimerization with ARNT and recruitment of CBP/p300 (*13–18*).

While this well-established canonical model explains important aspects of hypoxia-related pathophysiology, emerging clinical and mechanistic observations strongly suggest additional, non-traditional roles of HIF-2α (*3–5*, *9*, *19*, *20*). For example, pharmacologic inhibition of HIF-2α dimerization with HIF-2α-specific inhibitor Belzutifan is highly effective in VHL-mutant renal cancers but unexpectedly ineffective in neuroblastoma (*3*, *20*), indicating that key HIF-2α functions may operate outside the classical transcriptional paradigm.

To probe these non-canonical roles, we undertook a comprehensive structural and functional survey of the HIF-2α protein. Using an extensive deletion-construct library across two clinically relevant cancer models – neuroblastoma and clear cell renal cell carcinoma (ccRCC) – we mapped the specific domains required for controlling HIF-2α protein abundance, HRE-driven transcriptional activity, and cytoplasmic versus nuclear localization. Complementing this, we performed immunoprecipitation-mass spectrometry under both normoxic and hypoxic conditions, which enabled us to define the oxygen-dependent and oxygen-independent interactomes of HIF-2α.

Together, these analyses reveal a previously unrecognized level of structural and regulatory complexity within HIF-2α. These analyses identify domain functions that diverge strikingly from established paradigms and expose interaction networks not captured by classical models of hypoxia signaling. By uncovering regulatory determinants and binding partners that operate outside the traditional ARNT-dependent transcriptional framework, our work provides a fundamentally expanded view of HIF-2α biology; one that has direct implications for understanding its roles in tumor progression, metabolic regulation, and human disease.

## Results

### Generation of a HIF-2α deletion library to dissect domain-specific functions

The structural organization of HIF-2α is shown in Figure 1A. To investigate the determinants of its subcellular localization and transcriptional activity, we generated a library of HIF-2α variants cloned into the pcDNA3.1 vector with an N-terminal HA-tag. This library includes eleven individual deletion constructs of varying sizes that collectively span the entire coding sequence through overlapping regions (Fig. 1B). Additionally, we used two mutant constructs (Fig. 1B) to study nuclear translocation of HIF-2α: HIF-2α 2M, carrying proline-to-alanine substitutions at residues 405 and 531, which are key oxygen degradation domain (ODD) residues, and HIF-2α 3M, which includes an additional asparagine-to-alanine substitution at residue 847, important for FIH1 regulation (*17*, *21–23*).

**Figure 1.**
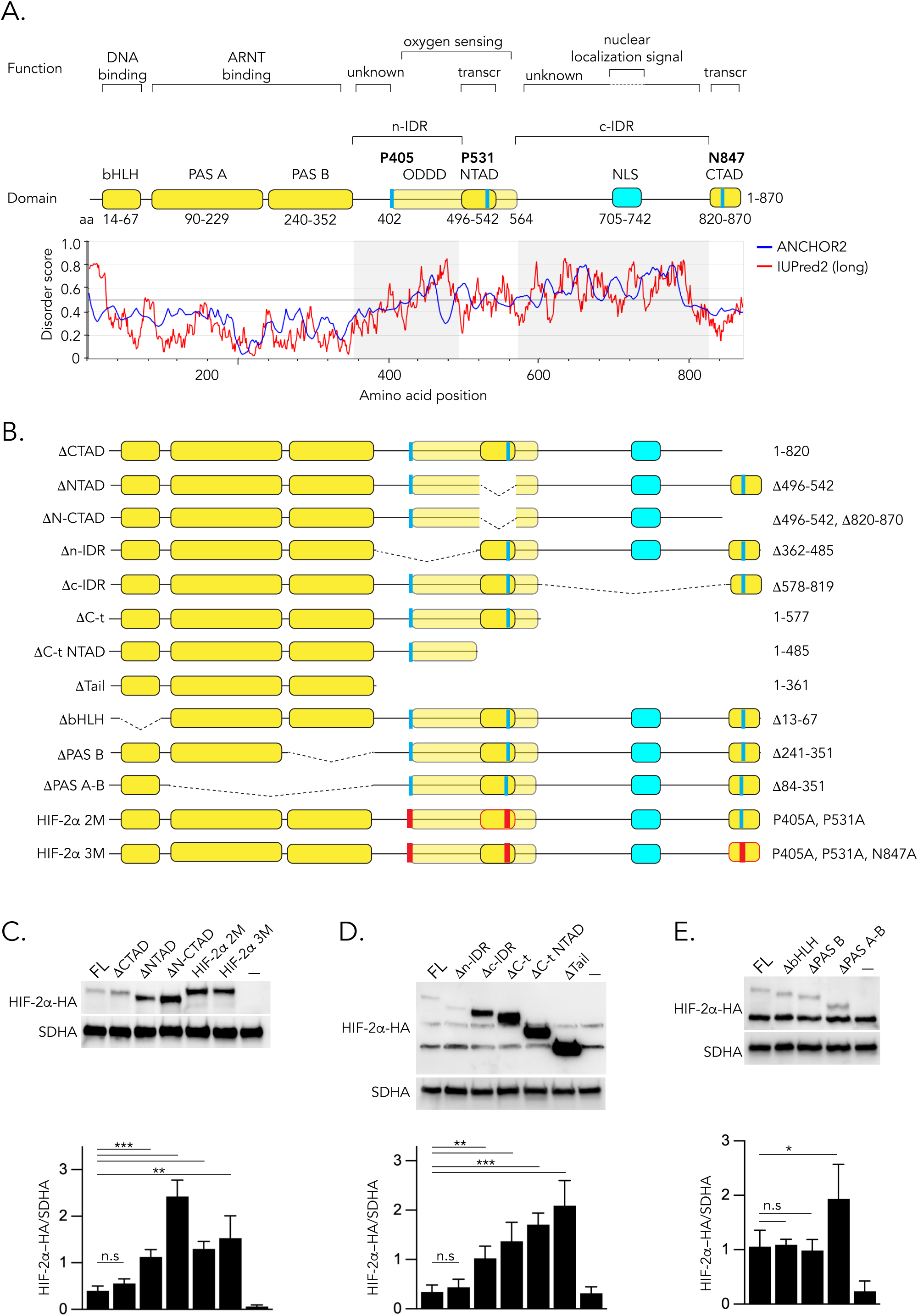
HIF-2α domains contribute differentially to protein abundance. (**A**) Schematic of the HIF-2α protein. Disorder prediction (IUPred2 (*43*)) identifies two intrinsically disordered regions (IDRs): n-IDR and c-IDR. bHLH, basic helix-loop-helix; PAS A/B, Per-Arnt-Sim domains; ODDD, oxygen-dependent degradation domain; NTAD/CTAD, N- and C-terminal transactivation domain; NLS, nuclear localization signal. (**B**) Overview of HIF-2α variants used in this study. Numbers indicate amino acid deletions or substitutions (substitutions shown in red). All constructs contain an N-terminal HA tag. (**C–E**) Expression of HIF-2α variants in SK-N-BE(2)c cells under normoxia (21% O2). Western blots were quantified, normalized to SDHA, and plotted from *n=4* independent experiments. Statistical analysis: unpaired two-tailed Student’s *t*-test (*n.s*, non-significant; * *p* < 0.05; ** *p* < 0.01; *** *p* < 0.001).

### PHD binding at P405 residue is not required for oxygen-dependent degradation of HIF-2α

We first assessed protein levels after transfection with our deletion and point-mutation variants from whole cell lysates of SK-N-BE(2)c neuroblastoma cells grown at normoxia (21% O₂; Fig. 1C–E), and found that the ΔNTAD, ΔN-CTAD, 2M, or 3M constructs resulted in markedly increased concentrations relative to full-length HIF-2α (Fig. 1C). This outcome is consistent with the loss of the major hydroxylation site P531, located within NTAD. Similarly, constructs with larger C-terminal deletions spanning these residues (ΔC-t, ΔC-t NTAD, and ΔTail) also exhibited elevated protein levels (Fig. 1D). In contrast, the Δn-IDR variant, despite lacking the P405 hydroxylation site, did not show increased expression (Fig. 1D). This indicates that removal of P405 alone is insufficient to stabilize HIF-2α and suggests that the P531 region has a greater influence on protein turnover. Deletion of domains unrelated to hydroxylation (ΔbHLH, ΔPAS-B, and ΔPAS-A-B) did not alter expression relative to full-length HIF-2α (Fig. 1E), consistent with retention of the principal oxygen-sensitive residues in these variants. Under hypoxic culture conditions (1% O₂; Supplementary Fig. S1A–C), expression levels for most variants mirrored those observed at normoxia.

Collectively, these results demonstrate that HIF-2α protein abundance is not governed solely by the presence or absence of individual hydroxylation sites. Instead, stability depends on the structural context of these sites, with the P531 residue playing a dominant role. Moreover, regulation of HIF-2α protein levels is at least partly independent of ambient oxygen concentration.

### The oxygen sensing domain of HIF-2α mediates its cytoplasmic localization

Examination of six neuroblastoma cell lines revealed consistent HIF-2α protein expression under normoxia (Fig. 2A). Subcellular fractionation of SK-N-BE(2) neuroblastoma cells cultured at normoxia or hypoxia revealed that HIF-1α is confined to the nuclear compartment (Fig. 2B–C). In contrast, HIF-2α was consistently detected within the cytoplasmic fraction across both oxygen conditions (Fig. 2B–C), in line with our previous observations in neuroblastoma and with clinical findings in paraganglioma (*3*, *4*).

**Figure 2.**
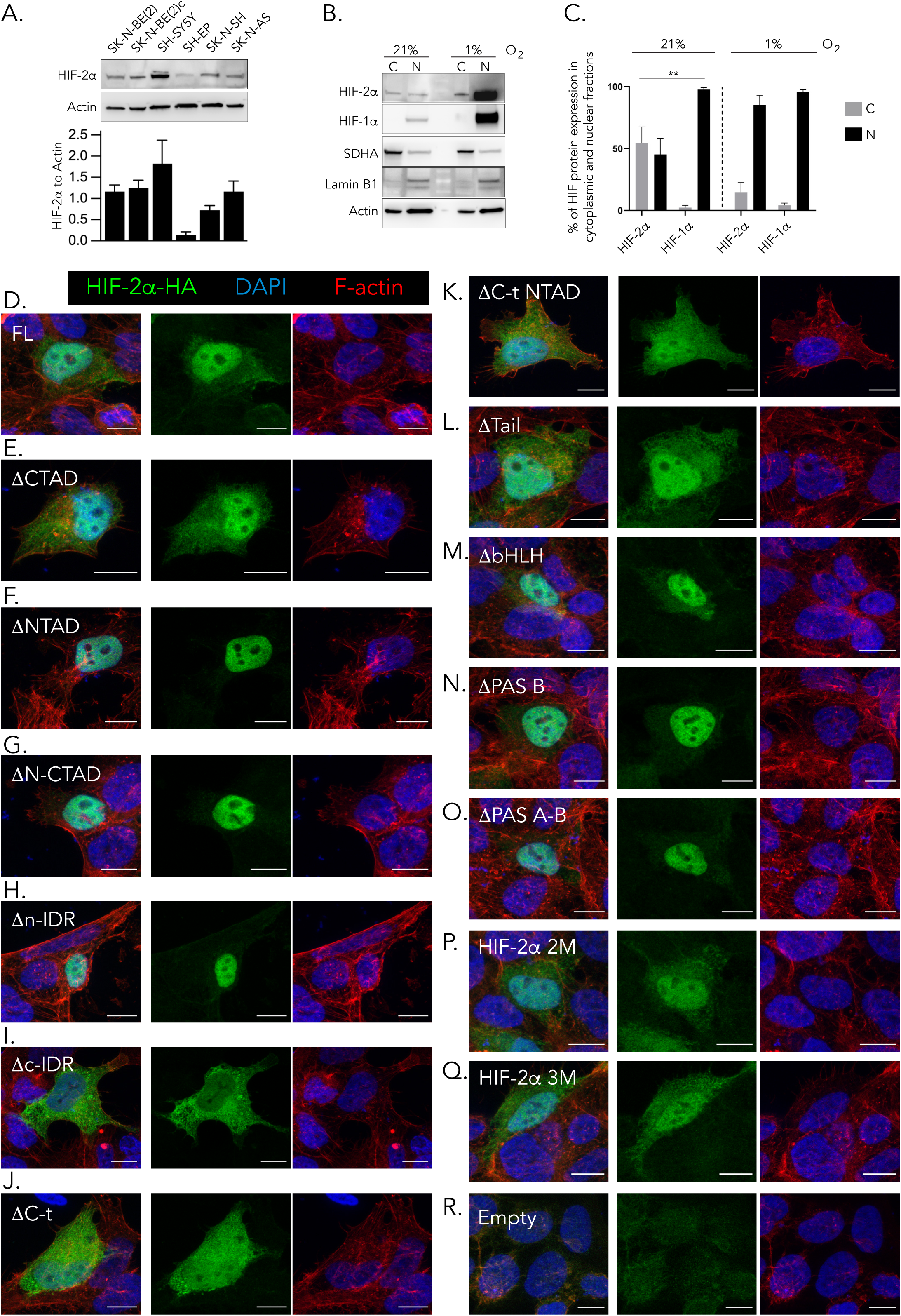
The oxygen-sensing domain of HIF-2α regulates its cytoplasmic localization. (**A**) Six neuroblastoma cell lines cultured at 21% O₂ were probed for endogenous HIF-2α. Western blot signals were quantified and normalized to Actin (loading control); *n*=3 independent experiments. (**B**) SK-N-BE(2) cells grown under normoxia (21% O₂) or hypoxia (1% O₂) were fractionated into cytoplasmic (C) and nuclear (N) compartments and analyzed for HIF-1α and HIF-2α. SDHA and Lamin B1 serve as cytoplasmic and nuclear markers, respectively. (**C**) Quantification of cytoplasm-to-nuclear ratios for HIF-1α and HIF-2α (*n*=3; unpaired two-tailed Student’s *t*-test; ** *p* < 0.01). (**D–R**) HA-tagged HIF-2α variants expressed in SK-N-BE(2)c cells were immunolabeled for the HA epitope; F-actin was counterstained with phalloidin, and nuclei visualized with DAPI. Scale bars: 10 µm. Representative images from *n*=6 independent experiments.

To delineate the domains governing HIF-2α subcellular localization, and thereby its non-canonical functions, we transiently expressed our 13 domain-altered HIF-2α variants (Fig. 1B) in SK-N-BE(2)c neuroblastoma cells and performed immunocytochemistry (Fig. 2D–R). Full-length HIF-2α localized to both the nucleus and cytoplasm (Fig. 2D), consistent with the fractionation data in Fig. 2B. Deletion of CTAD did not alter localization relative to full-length protein (Fig. 2E). By contrast, removal of NTAD, N-CTAD, or the n-IDR domains resulted in a marked loss of cytoplasmic signal with strong nuclear enrichment (Fig. 2F–H), indicating that the n-IDR and NTAD regions contribute to cytoplasmic retention. In the opposite direction, deletion of the c-IDR or the larger C-t region produced pronounced cytoplasmic accumulation (Fig. 2I–J), consistent with loss of the C-terminal nuclear localization signal (NLS) embedded within the c-IDR. Interestingly, deletion of NTAD or n-IDR in the context of the larger ΔC-t and Δc-IDR variants partially restored nuclear localization (Fig. 2K–L). These results suggest that elements within NTAD and n-IDR can counteract the absence of the C-terminal NLS, possibly by altering protein conformation or interaction networks that facilitate nuclear import. Removal of the bHLH, PAS-B, or combined PAS-A/B domains also reduced cytoplasmic localization (Fig. 2M–O), indicating that multiple structured regions cooperate to maintain a cytoplasmic pool of HIF-2α. In contrast, substitution of hydroxylation-associated residues at P405 and P531 (2M and 3M variants) did not alter the localization pattern (Fig. 2P–Q), demonstrating that perturbation of these individual sites within the ODD is insufficient to disrupt cytoplasmic distribution. Empty-vector transfection served as a negative control for HA staining (Fig. 2R).

We additionally examined full-length HIF-2α and selected variants (ΔNTAD, ΔN-CTAD, Δn-IDR, and Δc-IDR) in ccRCC 786-O and neuroblastoma SK-N-AS cells (Supplementary Fig. S2A–B, respectively), and found that across all three cell lines, the subcellular distribution of each variant mirrored the patterns observed in SK-N-BE(2)c cells. These results indicate that the localization rules encoded by these domains are conserved across human cell types.

Together, these data demonstrate that the c-IDR region, which contains one of two NLS, is essential for nuclear import of HIF-2α. This mirrors previous results for HIF-1α, in which the N-terminal basic region is not sufficient for nuclear localization in the intact protein, whereas the C-terminal NLS is required for nuclear import (*24*). Conversely, the PAS and ODD domains promote cytoplasmic localization under normoxic conditions. These domain-specific behaviors are maintained across cell types of different developmental origins, suggesting a conserved structural logic that dictates HIF-2α compartmentalization.

### HIF-2α NTAD domain is dispensable for HRE transactivation

We further transfected SK-N-BE(2)c neuroblastoma cells with the library of HIF-2α variants together with HRE-luciferase and CMV-Renilla reporters and performed dual-luciferase assays (Fig. 3A–D). Empty vector served as a negative control, and the 3M mutant, which lacks functional hydroxylation sites, served as a positive control. Treatment of the 3M variant with the HIF-2α-specific inhibitor PT2385/Belzutifan, which prevents ARNT binding, robustly suppressed reporter activity (Fig. 3C), confirming assay specificity. Deletion of the bHLH domain reduced HRE transactivation by ∼80% relative to full-length HIF-2α (Fig. 3B), consistent with the requirement for DNA binding. Removal of PAS-B or PAS-A/B domains decreased activity by ∼60% and ∼50%, respectively (Fig. 3B), highlighting the importance of ARNT heterodimerization for HIF-2α transcriptional activity. Deletion of the n-IDR, which includes the ODD, also reduced transactivation, whereas deletion of the c-IDR markedly enhanced it (Fig. 3C). C-terminal truncations revealed a graded requirement for the C-terminal region: ΔC-t reduced activity by ∼30%, whereas deletion of larger regions encompassing NTAD and CTAD (ΔC-t NTAD and ΔTail) completely abolished transactivation, despite elevated protein expression levels (Fig. 3D). Deletion of CTAD alone had no effect on HRE activation (Fig. 3A), whereas deletion of NTAD, individually or in combination with CTAD, increased transactivation. These results indicate that NTAD exerts a modulatory, rather than strictly activating, role in HIF-2α transcriptional activity.

**Figure 3.**
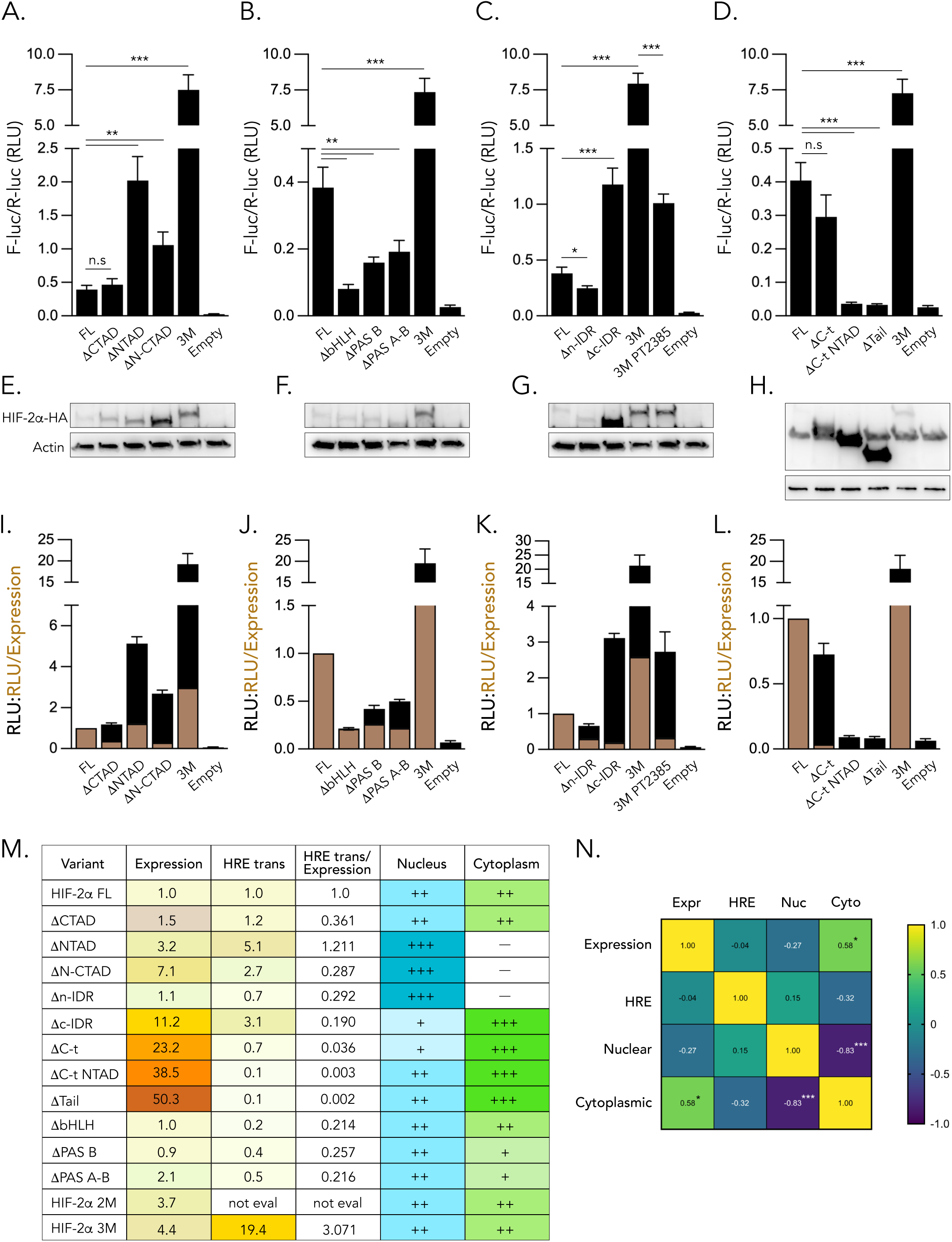
HIF-2α domains display independent functions. (**A–D**) HRE-luciferase reporter activity driven by HIF-2α variants. SK-N-BE(2)c cells were co-transfected with HIF-2α constructs, HRE-Luciferase, and CMV-Renilla, cultured at 21% O2, and assayed after 24 hours. Luciferase signals were normalized to Renilla and are shown as mean ± SD from *n*=3 independent experiments, each with three technical replicates (unpaired two-tailed Student’s t-test; * *p* < 0.05, ** *p* < 0.01, *** *p* < 0.001). FL, full-length. (**E–H**) A fourth technical replicate from samples analyzed in (A-D) was lysed for immunoblotting to verify construct expression. (**I-L**) Normalization of HIF-2α variants luciferase values to their protein expression. The black bars represent the same data from Fig. 3A-D that were normalized to full-length (FL) HIF-2α, with a set value of 1. Expression of the variants from 2 out of 3 luciferase experiments was quantified by densitometry, as detailed in the method section. The brown columns represent the relative luciferase signals normalized to relative HIF-2α variant expression. (**M**) Summary table showing expression, HRE transactivation, and subcellular localization of each HIF-2α variant at 21% O2. (**N**) Heatmap of Spearman correlation coefficients among HRE transactivation, protein expression, and nuclear and cytoplasmic localization. Color indicates the direction and strength of the correlation (Spearman’s r), with positive correlations shown in yellow/green and negative correlations in blue/purple. Asterisks indicate statistical significance (*n.s*, non-significant; * *p* < 0.05, *** *p* < 0.001).

To ensure that these differences did not arise from variable expression of the individual HIF-2α variants, we quantified protein concentration from whole cell lysates by Western blot (Fig. 3E–H) for each luciferase assay and normalized HRE transactivation of each HIF-2α variant to its relative protein levels (Fig. 3I–L). After accounting for protein abundance, deletion of CTAD or N-CTAD reduced transactivation, whereas ΔNTAD maintained slightly higher activity than full-length (Fig. 3I). Deletion of bHLH, PAS-B, PAS-A/B (Fig. 3J), n-IDR, or c-IDR (Fig. 3K) all decreased activity. As before, ΔC-t, ΔC-t NTAD, and ΔTail displayed virtually no transactivation (Fig. 3L).

A summary of the domain-specific effects on protein abundance, localization, and transactivation is shown in Fig. 3M. Correlation analyses revealed strong negative coupling between nuclear and cytoplasmic localization, and a modest positive correlation between cytoplasmic localization and overall protein abundance (Fig. 3N). Transcriptional activity did not correlate with either protein levels or subcellular localization (Fig. 3N), demonstrating that HIF-2α transactivation capacity is independent of protein concentrations or compartmental distribution.

### HIF-2α-driven tumor growth in vivo is partly cell subtype dependent

We xenografted SK-N-BE(2) and SK-N-AS neuroblastoma cells stably expressing either full-length HIF-2α or empty vector (Supplementary Fig. S3A–D) into the flanks of nude mice. Both cell lines express endogenous HIF-2α (Fig. 2A) and wild type VHL (*25*, *26*). In SK-N-BE(2) cells, enforced HIF-2α expression did not alter tumor formation rates or overall survival relative to controls (Fig. 4A–B). By contrast, SK-N-AS cells overexpressing HIF-2α exhibited a trend toward accelerated tumor onset and reduced early-stage survival compared with empty-vector controls (Fig. 4C–D), suggesting that tumor-promoting capacity of HIF-2α could be partly cell-subtype dependent.

**Figure 4.**
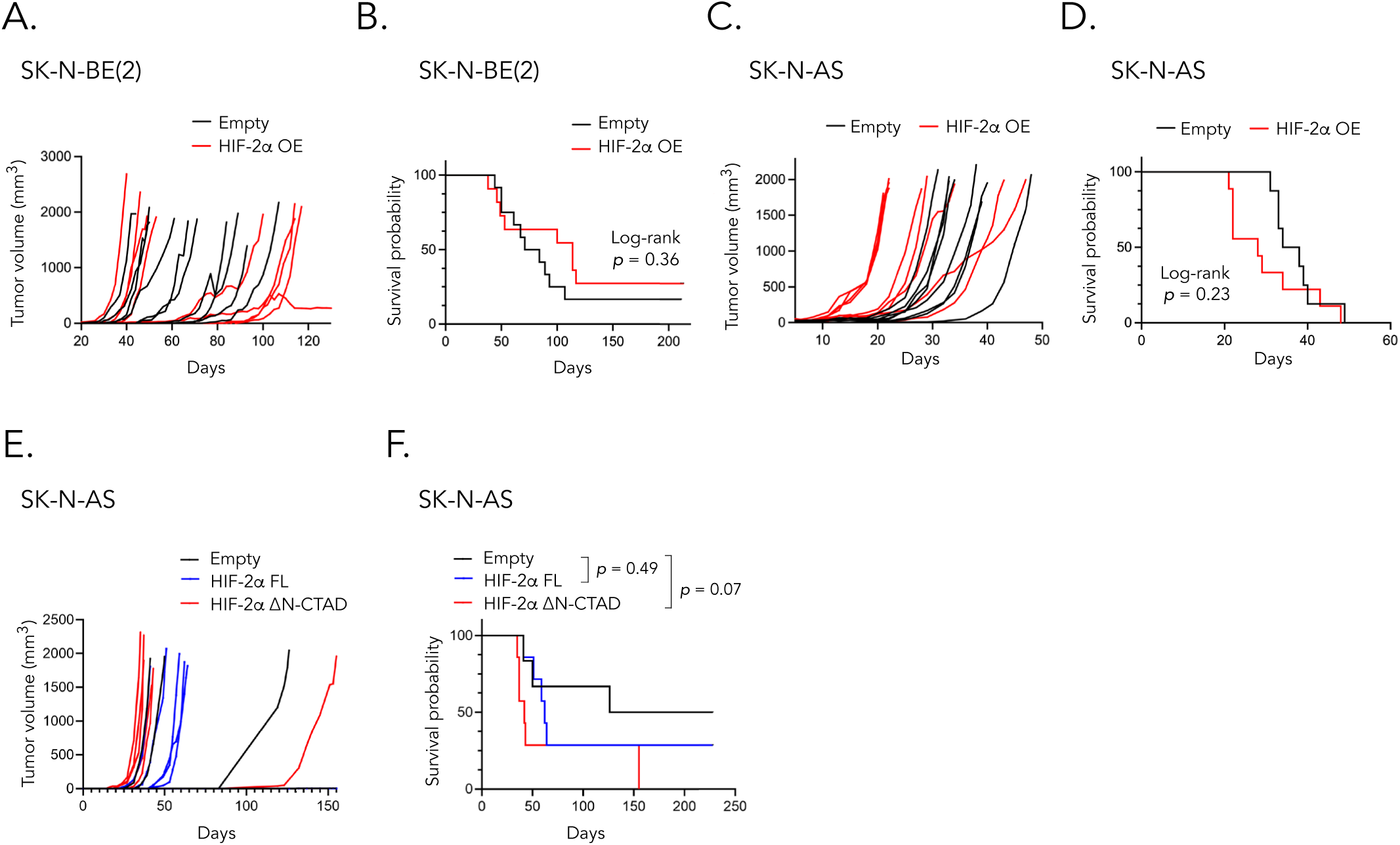
The transactivation domains of HIF-2α are dispensable for tumor growth in vivo. (**A–B**) Tumor growth and Kaplan-Meier survival of mice injected with neuroblastoma SK-N-BE(2) cells transduced with empty vector (EV) or HIF-2α overexpression (OE) constructs (*n*=10 EV; *n*=9 OE). (**C–D**) Tumor growth and survival in mice injected with neuroblastoma SK-N-AS cells transduced with empty vector or HIF-2α OE (*n*=8 EV; *n*=9 OE). (**E–F**) Tumor growth and survival in mice injected with SK-N-AS cells transfected with empty vector, HIF-2α full-length (FL), or ΔN-CTAD constructs (*n*=6 EV; *n*=7 FL; *n*=7 ΔN-CTAD). **(A-F)** Tumor volumes were measured every 2-3 days, and mice were euthanized when tumors exceeded 1800 mm³. Statistical significance between groups was determined by log-rank test.

### The transactivation domains of HIF-2α are dispensable for in vivo tumor growth

To determine whether the canonical transactivation domains contribute to HIF-2α-driven tumorigenesis, we injected SK-N-AS neuroblastoma cells transfected with full-length or ΔN-CTAD HIF-2α variants, or empty vector, subcutaneously into mice. Successful transfection was confirmed prior to implantation (Supplementary Fig. S3E–F). Tumor growth curves were similar across groups (Fig. 4E), but both full-length and ΔN-CTAD HIF-2α displayed a trend toward reduced survival relative to the empty-vector control (Fig. 4F), with the ΔN-CTAD variant showing the strongest effect. Tumor growth and survival did not differ between the full-length and ΔN-CTAD groups (Fig.4E–F).

These findings suggest that the N-terminal and C-terminal transactivation domains, classically required for ARNT-dependent HIF-2α transcriptional activity, are not necessary for HIF-2α to promote tumor progression *in vivo*. Instead, these results point to transactivation-independent, non-canonical mechanisms underlying HIF-2α-driven malignancy.

### Distinct HIF-2α structural variants elicit separable transcriptional signatures

RNA sequencing on wild type cells and cells expressing full-length HIF-2α, or the ΔN-CTAD and Δc-IDR variants, localizing the protein predominantly to the nucleus or cytoplasm, respectively, revealed that distinct structural features of HIF-2α differentially remodel the transcriptomic landscape. The *EPAS1/HIF2A* gene, encoding HIF-2α, was robustly upregulated across all transfected groups relative to wild type cells, confirming successful expression of the constructs (Fig. 5A–C). Differential expression analysis (padj < 0.05; log₂FC > 1) comparing full-length HIF-2α-expressing cells with wild type identified a similar number of upregulated and downregulated genes (10 and 8, respectively; Fig. 5A). Among the upregulated genes were *NDRG1* and *HIF1A*-*AS3*, both previously characterized as hypoxia-responsive, with prior studies further suggesting a potential functional relationship between these transcripts (*27–29*).

**Figure 5.**
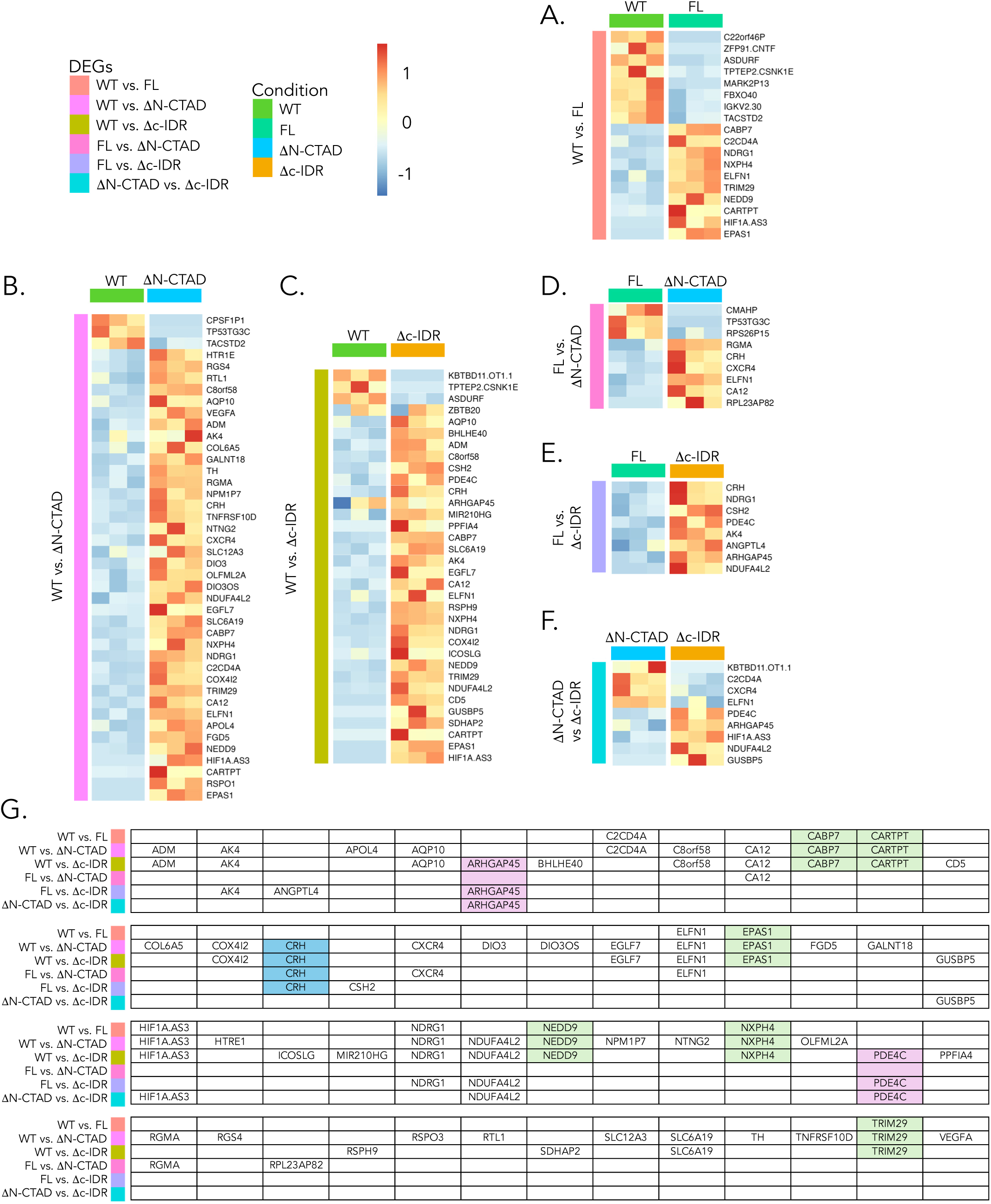
RNA sequencing reveals cytoplasmic- and nuclear-regulated genes. **(A–F)** RNA sequencing was performed on SK-N-BE(2) cells cultured for 48 hours at 21% O2 and either untransfected (wild type) or transfected with full-length (FL), ΔN-CTAD (driving nuclear localization), or Δc-IDR (driving cytoplasmic localization) HIF-2α constructs. Heatmaps of differentially expressed genes (DEGs) comparing: (**A**) wild type vs. HIF-2α FL; (**B**) wild type vs. ΔN-CTAD; (**C**) wild type vs. Δc-IDR; (**D**) FL vs. ΔN-CTAD; (**E**) FL vs. Δc-IDR; (**F**) ΔN-CTAD vs. Δc-IDR. **(G)** Summary of overlapping and unique DEGs across all comparisons. Genes in purple are upregulated exclusively by Δc-IDR relative to all other conditions; genes in green are upregulated by HIF-2α regardless of whether the construct is full-length or truncated; and the gene in blue is upregulated by either ΔN-CTAD or Δc-IDR but not by full-length HIF-2α, with no difference between the two deletion constructs.

We next stratified the dataset by performing all pairwise comparisons among the four conditions (wild type, full-length HIF-2α, ΔN-CTAD, and Δc-IDR) which yielded the final set of differentially expressed genes (DEGs; Fig. 5A–F). When comparing wild type cells with ΔN-CTAD-expressing cells (promoting nuclear localization), all genes upregulated by full-length HIF-2α were retained, and an additional 29 genes were uniquely induced, resulting in a total of 39 upregulated transcripts (Fig. 5B). Notably, several of these additional genes are well-established hypoxia-responsive targets, including *ADM*, *VEGFA*, *COX4I2*, *NDUFA4L2*, *CXCR4*, and *CA12*. In contrast, only three genes were downregulated in ΔN-CTAD cells, with just one overlapping with the full-length HIF-2α signature (Fig. 5B). A similar pattern emerged for the Δc-IDR variant (promoting cytoplasmic localization). Nine of the ten genes upregulated by full-length HIF-2α were also induced in Δc-IDR-expressing cells, together with 22 additional genes (31 total; Fig. 5C). Four genes were downregulated in Δc-IDR cells (Fig. 5C), two of which overlapped with those downregulated by full-length HIF-2α.

Mapping DEGs across all conditions revealed five genes, *CABP7*, *CARTPT*, *NEDD9*, *NXPH4*, and *TRIM29* (Fig. 5G; marked in green), that were consistently upregulated by HIF-2α, regardless of whether the protein was full-length or carried the ΔN-CTAD or Δc-IDR deletions. Their induction therefore appears independent of the transactivation domains, the c-IDR region, and the subcellular localization of HIF-2α (Fig. 5G). In contrast, *CRH* (marked in blue) was uniquely induced in both deletion-construct conditions but not by full-length HIF-2α, suggesting that its regulation does not depend on nuclear versus cytoplasmic localization *per se*, but instead requires the absence of either the transactivation domains or the c-IDR region (Fig. 5G). Additionally, *ARHGAP45* and *PDE4C* (marked in purple) were upregulated exclusively upon deletion of the c-IDR domain (Fig. 5G), implying that these transcripts may be preferentially influenced by cytoplasmic HIF-2α or that their induction is normally constrained by protein-protein interactions within the c-IDR.

### Global mapping of the HIF-2α interactome across oxygen conditions and tumor contexts

To define the protein interaction landscape of HIF-2α under distinct physiological conditions, we performed immunoprecipitation-mass spectrometry (IP-MS) screens (Fig. 6A) in neuroblastoma and ccRCC cells cultured at normoxia (21% O₂), as well as in neuroblastoma cells cultured at hypoxia (1% O₂). Hypoxic induction was validated by robust upregulation of HIF-1α, Dec1, and SerpinB9 (Supplementary Fig. S4A–B). Cells were transfected with HA- or FLAG-tagged HIF-2α expression constructs and subjected to affinity pulldown followed by liquid chromatography (LC)-MS/MS. A stringent filtering pipeline was applied to derive high-confidence interactors for each condition (Fig. 6A).

**Figure 6.**
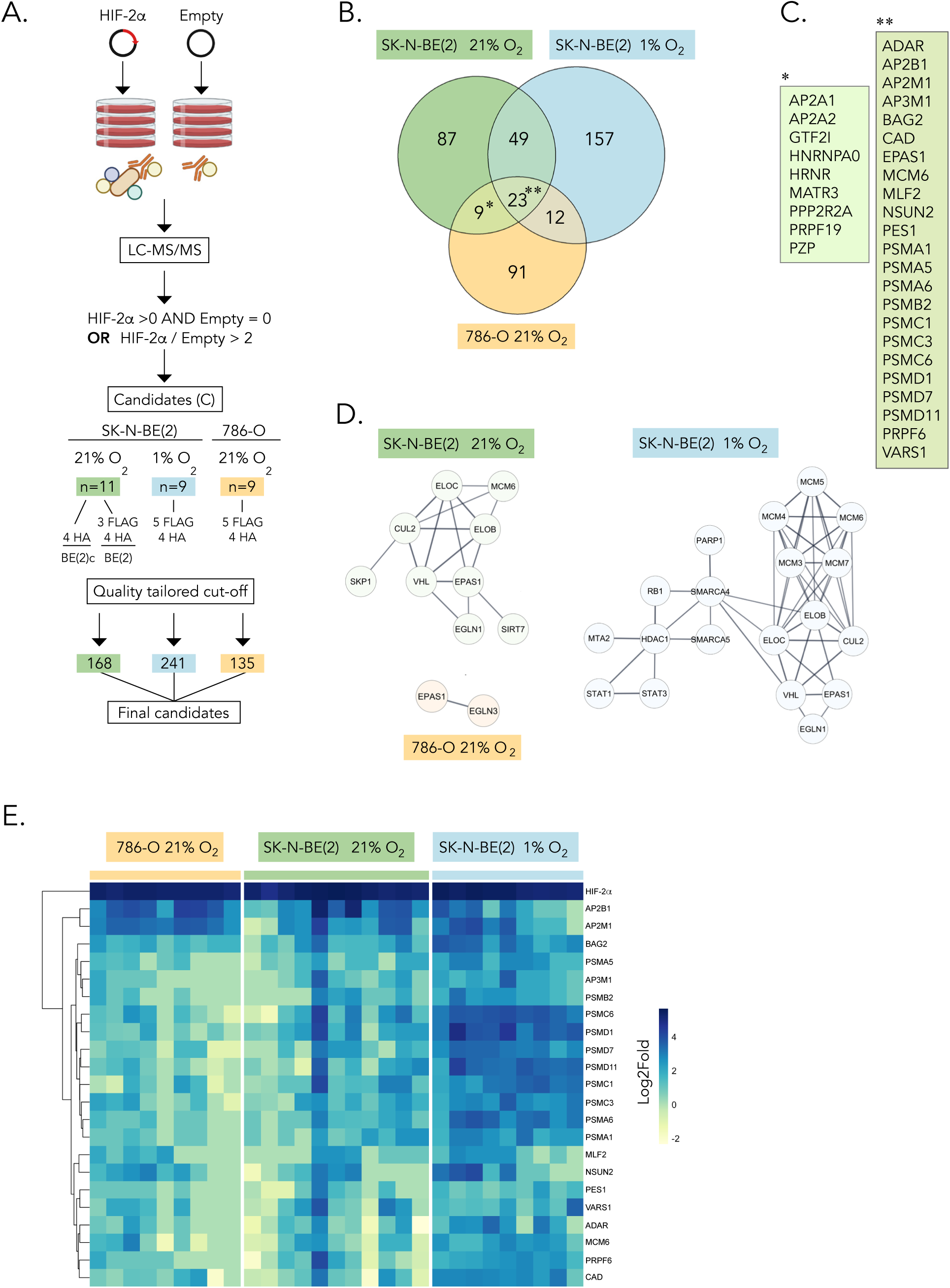
IP-MS screen identifies novel HIF-2α interactors. **(A)** Workflow schematic. Liquid Chromatography-tandem Mass Spectrometry (LC-MS/MS) data were processed with standard filtering; proteins were considered candidate interactors when candidate peptides were detected in HIF-2α but not Empty, or the HIF-2α:Empty peptide ratio exceeded 2. A total of *n*=11 experiments were performed in neuroblastoma SK-N-BE(2) cells at 21% O₂, *n*=9 at 1% O₂, and *n*=9 in renal cell carcinoma 786-O cells at 21% O2. (**B**) Venn diagram showing overlap of interactors among the three experimental groups. (**C**) List of shared interactors identified in (B). (**D**) STRING analysis of proximal HIF-2α interactors for each group, displaying high-confidence interactions (score ≥ 0.8). (**E**) Heatmap of log2fold-changes for the 23 common interactors comparing Empty vector vs. HIF-2α overexpression conditions.

Peptide-count correlation across replicates demonstrated high reproducibility (Supplementary Fig. S4C). HIF-2α itself was consistently detected in all samples, confirming successful enrichment of the bait protein. To generate final curated interactor lists within the top ranked 100–250 proteins per condition, thresholds were tailored to each dataset based on replicate correlation and overall data quality (Fig. S4C). Applying this filtering strategy yielded high-confidence interactomes comprising 135 proteins in ccRCC cells, and 168 proteins in normoxic and 241 in hypoxic neuroblastoma cells (Fig. 6A). Full datasets for all detected proteins are provided in Supplementary Table S1.

We next compared the three interactome datasets and identified 23 proteins shared across all conditions (Fig. 6B–C). In addition, 157 interaction partners were specific to hypoxia, while 9 proteins were consistently detected under normoxia in both neuroblastoma and ccRCC cells (Fig. 6B–C). To visualize known protein-protein interaction networks within these groups, we analyzed the curated interactor lists using STRING and Cytoscape (*30*, *31*). These analyses recovered well-established components of the HIF ubiquitin ligase machinery, ELOB, ELOC, VHL, CUL2, and EGLN1 (PHD2), as HIF-2α interactors in neuroblastoma cells under both normoxic and hypoxic conditions (Fig. 6D). As expected, ELOB, ELOC, VHL, and CUL2 were absent from the interactome in VHL-deficient 786-O cells, where HIF-2α instead associated with EGLN3/PHD3 (Fig. 6D). This observation is consistent with the requirement of VHL for recruitment of ELOB, ELOC, and CUL2 to assemble the HIF-α-specific ubiquitin ligase complex (*32*). Together, these results demonstrate that HIF-2α engages a conserved core of interactors across oxygen conditions and tumor contexts, while also recruiting oxygen-dependent and cell type-specific partners that likely contribute to its diverse cellular functions.

To determine whether the 23 shared interactors (Fig. 6C) exhibit differential abundance across conditions, we compared their log fold changes between empty vector controls and cells overexpressing full-length HIF-2α (Fig. 6E). As expected, HIF-2α itself was detected at high and comparable levels in all groups (Fig. 6E). Among the shared interactors, most proteins showed higher abundance under hypoxic conditions relative to normoxia, with the strongest shifts observed for proteasome-associated components (Fig. 6E). These data suggest that although these 23 proteins form a conserved core of the HIF-2α interactome, their relative abundance, and thus potentially their functional contribution, is partly modulated by oxygen availability.

### The HIF-2α interactome is enriched for nuclear transport and DNA-associated functions under hypoxia

Gene Ontology enrichment analysis of the cellular compartments, biological processes, and molecular functions associated with the interactomes identified under each condition (n = 241 for hypoxic neuroblastoma cells, n = 168 for normoxic neuroblastoma cells, and n = 135 for ccRCC cells) revealed a shared enrichment for proteasome- and ubiquitin-related pathways across all three groups (Fig. 7A–C; Supplementary Figs. S5–S7). This is consistent with the established role of ubiquitin-mediated degradation in HIF-2α regulation. Under hypoxic conditions, however, the HIF-2α interactome showed additional and striking enrichment for processes linked to nucleocytoplasmic transport, protein targeting to intracellular organelles, and regulation of DNA conformation (Fig. 7B). Correspondingly, several molecular functions associated with DNA handling, such as DNA helicase activity, were also represented (Fig. 7B).

**Figure 7.**
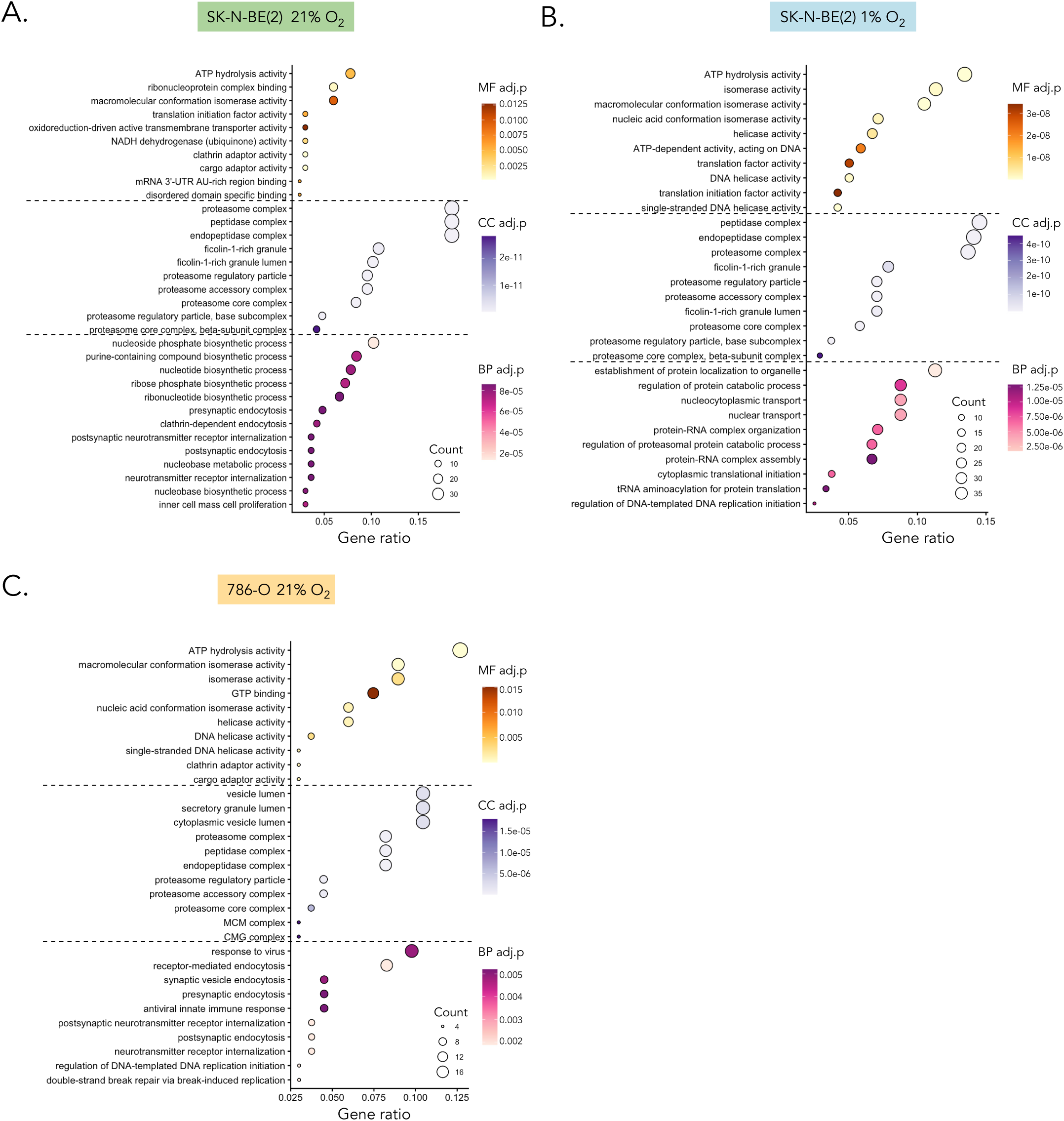
Gene ontology (GO) enrichment of HIF-2α interactors. **(A-C)** Dot plots show the top enriched GO terms ranked by significance. Dot size represents number of genes associated with each term, and dot color represents adjusted p-value with Benjamini-Hochberg correction. Only terms meeting significance threshold of *p* <0.05 were included. GO was performed for biological processes (BP), cellular component (CC), and Molecular function (MF). Neuroblastoma SK-N-BE(2) cells cultured in 1% oxygen **(A)** or 21% oxygen **(B)**, and renal cell carcinoma 786-O cells cultured in 21% oxygen **(C)**.

STRING network analysis further supported these findings, revealing interconnected clusters involving nuclear import machinery, DNA unwinding and replication factors, chromosome-maintenance proteins, and proteasome components (Supplementary Fig. S5). Proteins associated with mitochondrial, endoplasmic reticulum, and Golgi compartments were also detected within the hypoxic interactome (Supplementary Fig. S5). Together, these data indicate that under low-oxygen conditions, HIF-2α engages interaction networks that facilitate its nuclear import and interface with DNA-associated processes. This is consistent with the enhanced nuclear localization of HIF-2α observed during hypoxia and suggests that the hypoxic interactome actively supports both its subcellular trafficking and transcription-related functions.

### The normoxic HIF-2α interactome is enriched for translational and metabolic functions

In normoxic neuroblastoma cells, pathway analysis revealed enrichment of proteins involved in RNA synthesis and clathrin-mediated intracellular transport, including components of coated vesicle formation (Fig. 7A). The interactome also contained numerous factors associated with metabolic processes, particularly those linked to oxidoreductase activity and NADH metabolism (Fig. 7A). STRING network analysis reinforced these findings, highlighting large protein clusters associated with mitochondrial function and clathrin-dependent trafficking pathways (Supplementary Fig. S6). In line with these results and consistent with published observations that HIF-2α participates in translational regulation (*5*) the normoxic interactome included multiple translation-initiation factors (Supplementary Fig. S6). Together, our findings indicate that under normoxic conditions HIF-2α engages interaction networks involved in mitochondrial metabolism, RNA translation, and intracellular protein transport, supporting a role for cytoplasmic HIF-2α in coordinating metabolic and biosynthetic processes.

### The ccRCC HIF-2α interactome highlights intracellular transport and genome-associated functions

In VHL-mutant 786-O ccRCC cells, pathway analysis revealed a broader spectrum of enriched biological processes compared with neuroblastoma, including proteolysis, endocytosis, and DNA repair (Fig. 7C). The molecular functions represented within this interactome were similarly diverse, featuring GTPase activity and DNA helicase activity among others (Fig. 7C). Notably, however, as observed in normoxic neuroblastoma cells, HIF-2α also associated with clathrin-mediated intracellular transport machinery in ccRCC cells (Fig. 7B–C), indicating that this cytoplasmic trafficking pathway is a conserved feature of the HIF-2α interactome across tumor types. STRING network analysis further supported these findings, identifying vesicular transport modules and DNA replication/repair complexes, together with ribosomal proteins, as the most prominent interaction clusters (Supplementary Fig. S7).

### Core components of the HIF-2α interactome form interconnected protein complexes

To further dissect the organizational architecture of the HIF-2α interactome, we performed STRING analysis on the nine normoxia-specific cell type-independent interactors and the 23 proteins shared across all three datasets (Fig. 8A–B). Among the nine cell type-independent proteins, two members of the AP-2 family interconnected (Fig. 8A). As expected, within the 23 commonly detected interactors, proteasome-associated components formed a highly interconnected module (Fig. 8B).

**Figure 8.**
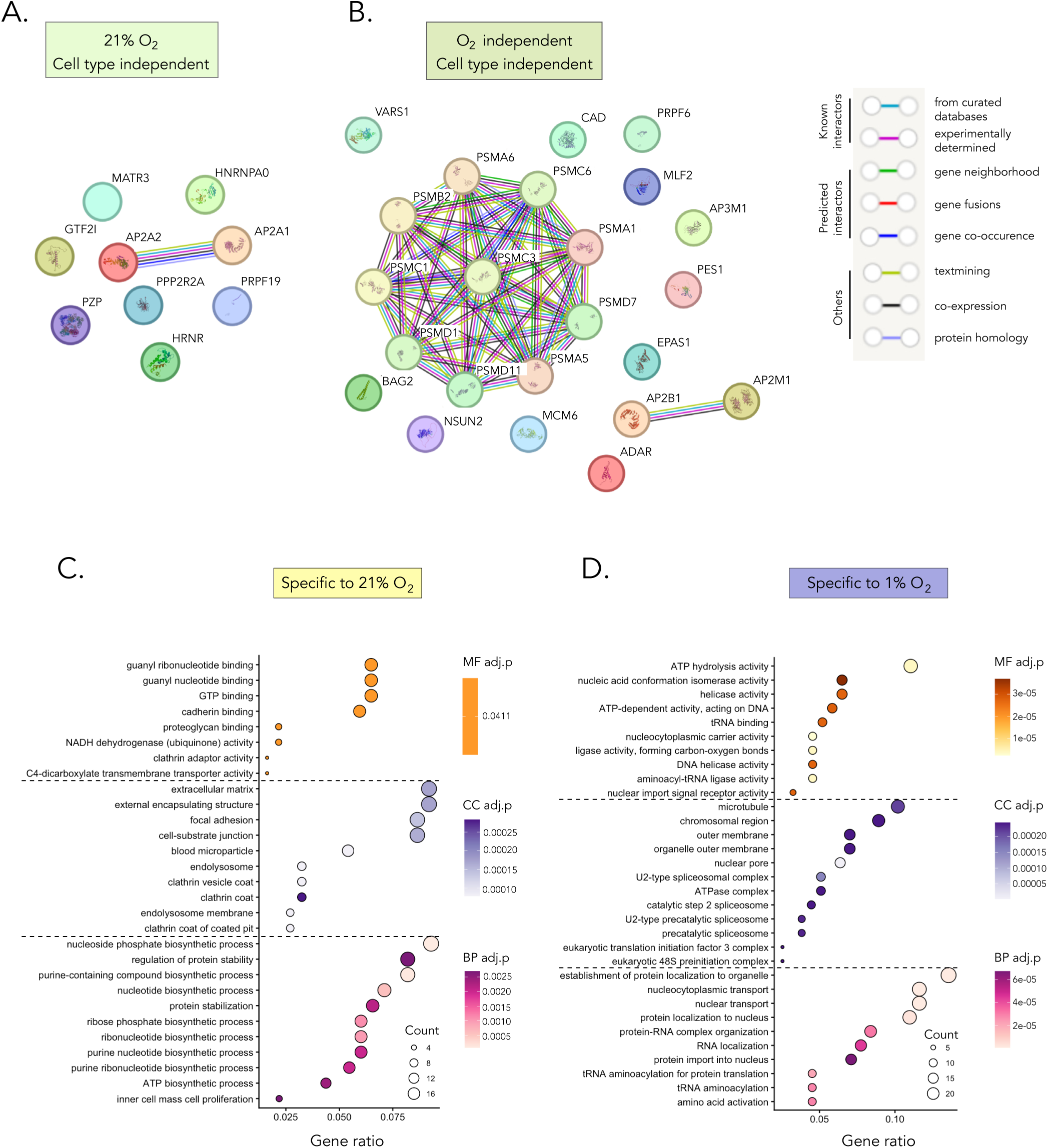
HIF-2α interactors are associated with distinct processes in different settings. **(A)** STRING analysis of proteins identified as normoxic-specific cell type-independent (proteins listed in Fig. 6C). **(B)** STRING analysis of proteins identified as oxygen- and cell type-independent (proteins listed in Fig. 6C). **(C-D)** Gene ontology (GO) enrichment analysis performed for biological processes (BP), cellular component (CC), and Molecular function (MF). Dot plots show the top enriched GO terms ranked by significance. Dot size represents number of genes associated with each term, and dot color represents adjusted p-value with Benjamini-Hochberg correction. Only terms meeting significance threshold of *p* <0.05 were included. Proteins detected only in neuroblastoma SK-N-BE(2) and renal cell carcinoma 786-O cells cultured in 21% oxygen **(C)**, and proteins detected only in neuroblastoma SK-N-BE(2) cells cultured in 1% oxygen **(D)** were analyzed.

### Normoxic HIF-2α associates with metabolic and mitochondrial functions

We separately analyzed proteins specific to normoxia (overlapping in neuroblastoma and ccRCC, i.e., cell-type independent; n = 187) and hypoxia (neuroblastoma; n = 157). Gene ontology enrichment of the normoxic interactors revealed strong overrepresentation of metabolic and mitochondrial processes, including translation, ATP and ribose-phosphate biosynthesis, and clathrin-mediated vesicle formation (Fig. 8C). These findings align with the broader interactome analyses demonstrating that normoxic HIF-2α engages cytoplasmic pathways involved in mitochondrial activity, protein synthesis, and intracellular trafficking. In contrast, hypoxia-specific interactors were enriched for pathways associated with cellular localization and nuclear transport and mapped to chromosomal and spliceosomal complexes (Fig. 8D). These functions are consistent with heightened nuclear engagement of HIF-2α at low oxygen levels, where transcriptional regulation dominates its activity. Together, these oxygen-stratified interaction profiles support a model in which normoxic HIF-2α participates primarily in metabolic and translational programs, while hypoxic HIF-2α interfaces with nuclear transport and chromatin-associated processes.

### HIF-2α localizes to mitochondria, ribosomes, and the endoplasmic reticulum

Given the enrichment of organelle-associated interactors and pathways in our IP-MS analyses, we next examined the subcellular distribution of endogenous HIF-2α by co-staining SK-N-BE(2) cells cultured at 21% O2 with markers for mitochondria (TOMM20), ribosomes (RPS6), the endoplasmic reticulum (P4HB), the Golgi apparatus (Gorasp2), and the cytoskeleton (phalloidin/F-actin). Immunofluorescent imaging (Fig. 9A–E) and quantitative co-localization analysis (Fig. 9F) demonstrated clear overlap between HIF-2α and mitochondrial structures (Fig. 9B), consistent with the identification of mitochondrial proteins and import machinery in the normoxic interactome. HIF-2α also co-localized with ribosomal (Fig. 9A) and ER-associated proteins (Fig. 9C), supporting a role for cytoplasmic HIF-2α in translational and biosynthetic processes. In contrast, we detected no appreciable co-localization with Golgi markers (Fig. 9D) or the actin cytoskeleton (Fig. 9E).

**Figure 9.**
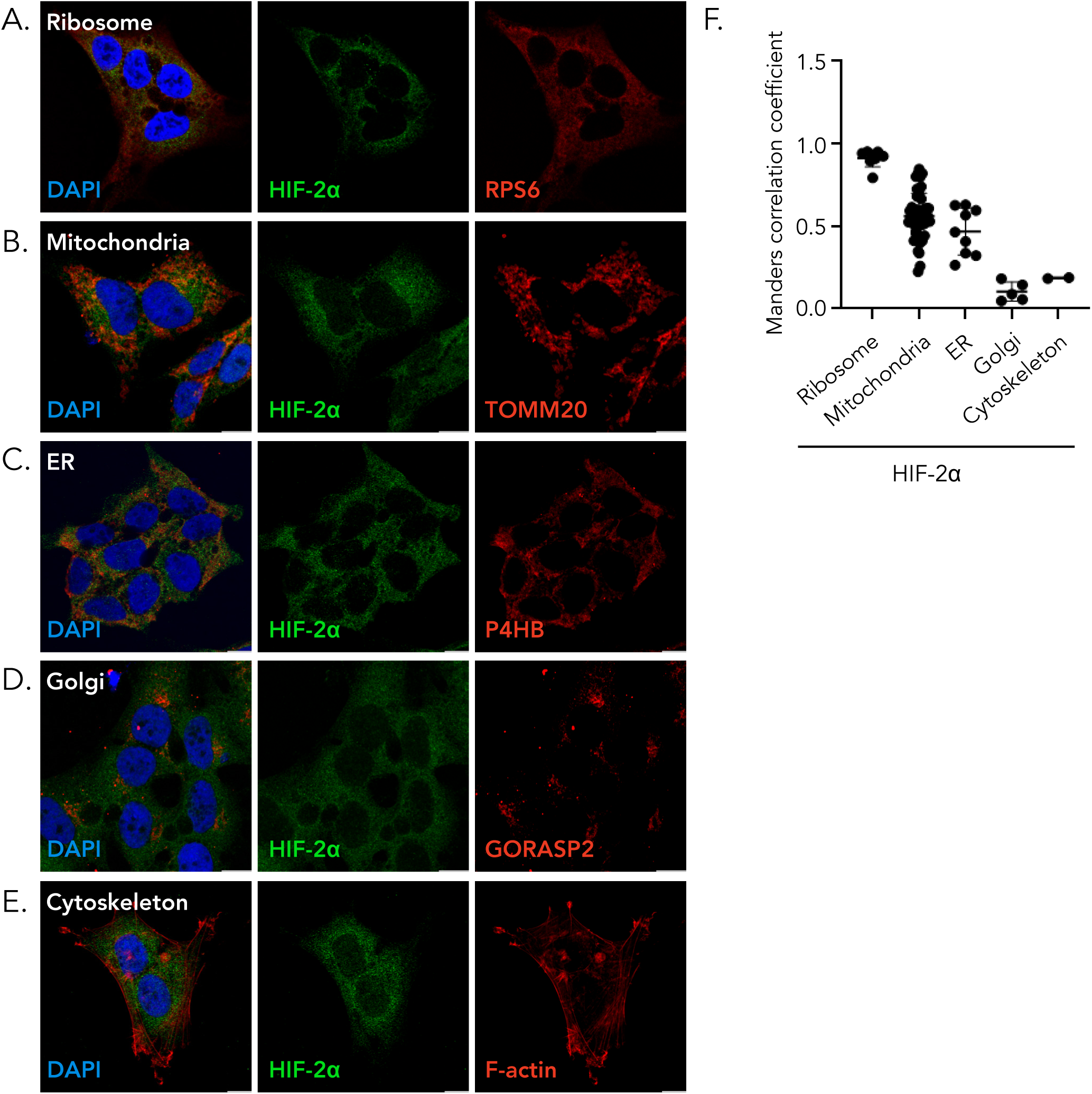
Endogenous HIF-2α colocalizes with ribosome and mitochondria. **(A-E)** Immunocytochemical staining of endogenous HIF-2α and ribosomal (RPS6; **A**), mitochondrial (TOMM20; **B**), Endoplasmic Reticulum (ER) (P4HB; **C**), Golgi (GORASP2; **D**), and cytoskeleton (F-actin; **E**) markers in SK-N-BE(2)c cells cultured at 21% O2. Scale bars, 10 µm, *n*=2 independent experiments. **(F)** Co-localization between cytoplasmic HIF-2α and individual subcellular markers was quantified using the Manders correlation coefficient. Each dot represents one measured cell. Horizontal bars indicate mean ± SD. Higher Manders correlation coefficients indicate greater spatial overlap between cytoplasmic HIF-2α signal and the indicated marker.

## Discussion

We provide a comprehensive characterization of how distinct subdomains of HIF-2α govern protein stability, transactivation capacity, and both canonical and non-canonical patterns of subcellular localization. In parallel, we delineate the HIF-2α interactome, offering an integrated view of its regulatory landscape. To contextualize these findings, we studied cells from two complementary tumor types in which HIF-2α contributes to disease progression through mechanistically different routes: RCC cells lacking VHL, where HIF-2α functions as a *bona fide* oncogenic driver, and neuroblastoma cells, in which non-canonical, oxygen-independent HIF-2α localization in well-oxygenated niches is strongly associated with poor prognosis and metastatic dissemination.

HIF-2α has emerged as an important therapeutic target due to its oncogenic roles across several tumor types, with FDA-approved inhibitors now in clinical use for RCC and late-stage clinical evaluation in PPGL. Small-molecule screening identified Belzutifan (PT2385/PT2399) as a HIF-2α-specific antagonist that binds a unique pocket within the PAS-B domain, sterically preventing ARNT heterodimerization and thereby blocking HIF-2-dependent transcription (*33*). Although Belzutifan confers substantial clinical benefit in a subset of patients with RCC as demonstrated in Phase III clinical trials (e.g., NCT04195750) (*34*), data from Phase I studies (NCT02293980) (*35*), together with preclinical analyses in ccRCC cell lines and patient-derived xenograft models (*36*, *37*), reveal that both intrinsic and acquired resistance are common. Many tumors fail to respond from the outset, while others develop resistance over the course of treatment. In a subset of cases, prolonged treatment failure has been linked to acquired mutations in HIF-2α that impair drug binding or downstream inhibition (*35*). However, these alterations only account for a fraction of non-responders, suggesting that additional undefined mechanisms contribute to resistance in the remaining non-responders. Notably, we previously demonstrated that neuroblastoma cells are largely resistant to Belzutifan both *in vitro* and *in vivo* (*3*). The cytoplasmic, oxygen-independent stabilization of HIF-2α in these cells, together with their preserved tumor-promoting behavior despite pharmacologic suppression of HIF-2-driven transcription strongly suggests that HIF-2α exerts critical non-canonical, non-transcriptional functions. These findings underscore the need to define the full spectrum of HIF-2α activities across cancer types to avoid ineffective treatment strategies and unnecessary adverse effects.

To elucidate the non-canonical mechanisms of HIF-2α and determine which protein domains regulate its distinct functions, we generated a library of HIF-2α variants and analyzed their behavior in our HIF-2α-dependent tumor cell models. Our data reveal that the region surrounding the P531 residue is critical for HIF-2α degradation, whereas the regions encompassing P405 and N847 do not exert comparable effects. The N847 residue, located within the CTAD, is the FIH hydroxylation site that sterically interferes with co-activator binding (*12*, *13*). In contrast, P405 and P531 are prolyl hydroxylation sites targeted by PHDs, marking HIF-2α for VHL-mediated ubiquitination (*22*). Notably, these two residues reside in distinct structural environments within the ODD: P531 lies within the NTAD, whereas P405 is positioned in the adjacent n-IDR. Both P405 and P531 support PHD-dependent VHL recruitment; however, our findings indicate that P531 plays a disproportionately important role in regulating HIF-2α stability. This observation aligns with clinical evidence where missense mutations affecting P531 and nearby residues within *EPAS1* constitute well-established pathogenic hotspots in pheochromocytoma, paraganglioma, and erythrocytosis (*19*, *38–40*). These mutations impair hydroxylation at P531, resulting in increased HIF-2α stability and prolonged protein half-life. Together, these data underscore the central role of the P531 region in controlling HIF-2α turnover and disease-associated dysregulation.

We found that deletion of the C-terminal IDR (Δc-IDR) results in a striking shift of HIF-2α toward the cytoplasm, with nuclear expression almost completely abolished. This phenotype is consistent with the removal of the C-terminal NLS located within this region. Notably, the Δc-IDR construct retains the second NLS at the N-terminus; however, our data show that this N-terminal motif is insufficient to mediate nuclear import of full-length HIF-2α. This observation aligns with previous findings for HIF-1α, where the N-terminal basic region does not function as an NLS in the context of the intact protein, and nuclear import instead requires the C-terminal NLS (*24*). Importantly, none of the other deletion constructs, those retaining the c-IDR region, display reduced nuclear localization, underscoring the essential role of the c-IDR in nuclear import of HIF-2α. Because the Δc-IDR construct eliminates the full region rather than selectively removing the C-terminal NLS, it remains possible that additional non-canonical nuclear localization determinants within the c-IDR contribute to nuclear shuttling, potentially via protein-protein interactions. Collectively, these findings highlight the c-IDR as a critical determinant of HIF-2α nuclear localization.

Interestingly, although deletion of the c-IDR alone redirects HIF-2α to the cytoplasm, the additional removal of CTAD (ΔC-t), CTAD together with NTAD (ΔC-t NTAD), or CTAD, NTAD, and the n-IDR (ΔTail) progressively attenuates this effect. In other words, the more extensive the C-terminal deletions become, the less pronounced the cytoplasmic accumulation appears. The mechanism underlying this unexpected pattern remains to be determined, but one possible explanation is that domains outside the c-IDR confer alternative structural or protein-interaction cues that override the loss of the C-terminal NLS. For example, deletion of larger regions may disrupt binding interfaces required for conformational states that favor cytoplasmic retention or alternatively may remove interaction motifs that normally facilitate nuclear exclusion when the c-IDR is absent. Another possibility is that larger C-terminal deletions unmask the cryptic N-terminal NLS, which has been shown to be functional outside the context of the full-length protein in previous studies as discussed above (*24*).

The mechanisms governing cytoplasmic localization of HIF-2α appear to be more complex than the presence or absence of the C-terminal IDR alone. Deletion of NTAD, either alone or together with CTAD, results in nuclear import. In contrast, deletion of CTAD alone has no measurable effect on subcellular distribution, indicating that NTAD, but not CTAD, contributes to cytoplasmic retention of HIF-2α. These findings further suggest that transactivation capacity itself is not the determinant of HIF-2α localization. Moreover, removal of the n-IDR also drives nuclear import. Because both NTAD and n-IDR reside within, or encompass, the ODD, these observations support the idea that oxygen-sensing elements within the ODD participate in restoring HIF-2α into dual cytoplasmic and nuclear expression. However, when NTAD and n-IDR are deleted in combination with the c-IDR (ΔTail), the phenotype reverses: the protein localizes also to the cytoplasm and exhibits elevated levels. Thus, the C-terminal NLS within the c-IDR appears sufficient to drive nuclear import, consistent with the Δc-IDR phenotype. Yet, because full-length HIF-2α normally localizes to both the cytoplasm and nucleus under physioxic and hypoxic conditions, this NLS alone is not enough to ensure exclusive nuclear localization. We therefore propose that subcellular distribution of HIF-2α emerges from an interplay between the C-terminal NLS, oxygen-dependent regulation mediated through the ODD, and interactions with additional protein partners that bind HIF-2α within this region. This combinatorial mechanism likely fine-tunes HIF-2α localization in response to cellular context and oxygen availability.

Substitution of P405 and P531, despite their positions within the n-IDR and NTAD, respectively, does not alter HIF-2α localization, indicating that these residues alone are insufficient to mediate cytoplasmic retention. Although the PAS domains of HIF-2α are essential for ARNT heterodimerization, it remains unclear whether this complex forms in the cytoplasm and contributes to nuclear import. Our results shed light on this question: deletion of PAS-B, and particularly deletion of both PAS-A and PAS-B, largely abolishes cytoplasmic localization. This pattern argues that ARNT-dependent dimerization is not required for nuclear import of HIF-2α and that the PAS domains instead participate in maintaining a cytoplasmic pool, potentially through structural constraints or protein-interaction interfaces independent of ARNT.

Despite unchanged subcellular localization following bHLH deletion and reduced cytoplasmic localization upon deletion of the PAS domains, which mediate DNA binding and ARNT dimerization, respectively, both variants exhibit markedly reduced transcriptional activity. Thus, nuclear presence alone is insufficient for full HIF-2α transactivation; the ability to form functional transcriptional complexes and engage target DNA is essential. Notably, removal of the bHLH or PAS domains does however not completely abolish transcriptional output, indicating that additional regions of HIF-2α contribute to basal or compensatory transactivation capacity. In contrast, deletion of the entire C-terminal region, comprising NTAD, c-IDR, and CTAD (ΔC-t NTAD), or deletion of this region together with the n-IDR (ΔTail) eliminates transactivation altogether. These findings underscore that the combined action of NTAD, c-IDR, CTAD, and the adjacent n-IDR is indispensable for transcriptional competence. Given this, one might expect the classical transactivation domains, NTAD and CTAD, to be strictly required for transcription. However, our results reveal a more nuanced regulatory logic. Deletion of NTAD alone paradoxically increases transcriptional activity, suggesting that this domain exerts a partial inhibitory or modulatory function under basal conditions. Deletion of CTAD alone has no effect on transactivation, and when CTAD removal is combined with NTAD deletion, it partially counteracts the transcriptional increase observed in the ΔNTAD variant. These patterns indicate that NTAD and CTAD do not function simply as additive activators; rather, they appear to cooperate in a context-dependent manner, with NTAD exerting a restraining influence that can be unmasked when CTAD remains intact. Together, these data highlight an unexpected complexity in the architecture of the transactivation machinery of HIF-2α.

While deletion of CTAD does not affect subcellular localization or transactivation potential, removal of NTAD leads to complete nuclear localization. A previous study demonstrated that NTAD alone is sufficient to drive HRE transactivation in the absence of CTAD in U2OS and 786-O cells (*41*). An important methodological difference is that Yan and colleagues used P405A/P531A and P405A/P531A/N847A backgrounds, corresponding to our HIF-2α 2M and 3M variants, for their deletion constructs, whereas our experiments were performed in the context of full-length HIF-2α. Nonetheless, our findings converge in showing that combined deletion of NTAD and CTAD does not fully abolish HRE promoter activation (*41*), suggesting the presence of an additional, cryptic transactivation site. Our deletion of the n-IDR results in reduced transactivation, which could be attributed either to a steric effect on the PAS-B domain, potentially altering the spatial relationship to NTAD, or to the removal of this putative cryptic transactivation element. Notably, Yan et al. did not evaluate subcellular localization of their variants, leaving open the possibility that differences in localization dynamics may also contribute to the observed functional discrepancies.

Adding yet another layer of complexity to the regulation of HIF-2α transactivation, deletion of the c-IDR, which contains the C-terminal NLS but has otherwise poorly defined functions, paradoxically increases transcriptional activity despite driving the protein almost entirely to the cytoplasm. Although this could theoretically stem from a small residual nuclear pool of HIF-2α, a more compelling interpretation is that cytoplasmic HIF-2α itself can support transcriptional output through ‘transcriptional islands’ present throughout the cell. This interpretation is reinforced by our previous observations in neuroblastoma cells, where treatment with PT2385 abolishes HIF-2α-ARNT dimerization and increases cytoplasmic localization, yet downstream gene expression remains largely unchanged (*3*). In contrast, knockdown of HIF-2α markedly reduced target gene expression, indicating that transcriptional programs attributed to HIF-2α persist even when its nuclear functions are compromised, and supporting the existence of non-canonical, non-nuclear roles for HIF-2α in the cytoplasm (*3*). Together, these results demonstrate that nuclear localization *per se* does not dictate the transcriptional output of HIF-2α. They also reveal that targeting individual transactivation domains is insufficient to suppress HIF-2α signaling, underscoring the importance of considering both canonical and non-canonical regulatory mechanisms when designing therapeutic strategies.

While CTAD is dispensable for HIF-2α function, NTAD emerges as a key regulator of HIF-2α homeostasis. The role of NTAD in regulating subcellular localization, transactivation and protein abundance is notably complex. Deletion of NTAD increases HIF-2α protein levels, and this effect is further amplified when NTAD is removed together with the c-IDR. Yet the combined deletion fundamentally alters the phenotype: whereas NTAD deletion alone promotes nuclear localization and enhanced transactivation, simultaneous removal of the c-IDR instead drives cytoplasmic localization and reduces transactivation. These contrasting outcomes suggest that protein-protein interactions with HIF-2α are highly sequence- and context-dependent. We propose that partner proteins binding specifically to NTAD recruit a distinct set of interacting factors, generating unique complexes or steric constraints that influence nuclear import and transcriptional activity of HIF-2α. In contrast, interactions occurring at the NTAD-c-IDR interface, or within c-IDR alone, likely engage an alternative cohort of partner proteins or induce different conformational states that can amplify, shift, or even counteract the effects mediated by NTAD alone. The observation that a domain traditionally associated with transcriptional activation also regulates protein stability and subcellular distribution is previously unrecognized and underscores that HIF-2α regulation is far more intricate than appreciated to date. Identifying the specific proteins that interact with NTAD, c-IDR, and their interface, and dissecting how these interactions cooperate or compete, will be crucial to fully understand the tumor-promoting functions of HIF-2α.

Strikingly, neither the C-terminal nor the N-terminal transactivation domains are required for tumor growth *in vivo*, revealing that the tumor-promoting activity of HIF-2α is, at least in part, mediated through mechanisms that extend beyond its canonical role in driving transcription of downstream target genes. This is an unexpected and conceptually transformative finding. It challenges the long-standing assumption that HIF-2-dependent tumorigenesis is primarily explained by transcriptional activation of hypoxia-responsive programs. Instead, it places renewed emphasis on non-canonical, non-nuclear functions of HIF-2α, functions that may involve cytoplasmic signaling events, protein-protein interactions, or scaffolding roles that shape tumor cell behavior independently of ARNT dimerization and DNA binding. In the context of our broader domain-mapping data, these results underscore that HIF-2α possesses oncogenic capacities that are uncoupled from its classical transactivation machinery, necessitating a fundamental re-evaluation of how this protein fully drives malignancy.

Using IP-MS, we mapped the HIF-2α interactome, identifying both individual protein partners and the broader pathways in which these complexes participate. Proteins that associate with HIF-2α under hypoxic conditions predominantly function in nuclear import and DNA-binding processes, consistent with canonical transcriptional regulation. In contrast, proteins binding to normoxic HIF-2α are enriched in pathways related to mitochondrial metabolism, protein translation, and subcellular trafficking. Supporting these biochemical data, we observe that cytoplasmic HIF-2α co-localizes with ribosome, mitochondria and endoplasmic reticulum. Together, these findings suggest that the tumor-promoting functions of cytoplasmic HIF-2α are driven, at least in part, by its engagement in metabolic and translational processes rather than by classical transcriptional activity. This shift in functional context aligns with our broader domain-mapping results and further reinforces the emerging view that HIF-2α possesses powerful non-canonical roles.

In conclusion, we show that the processes regulating HIF-2α protein abundance, transcriptional activity, and subcellular localization operate at least partly independently, indicating that HIF-2α is controlled by multiple, distinct regulatory layers. We therefore hypothesize that additional unidentified parameters play crucial roles in shaping both the behavior of the HIF-2α protein itself and its diverse effects on cellular function. These insights have important pre-clinical and clinical implications. Many studies aimed at defining HIF-2α function have been guided by early foundational work emphasizing its nuclear localization and oxygen-dependent transcriptional activity. However, our findings highlight a greater level of regulatory complexity, indicating that cytoplasmic functions of HIF-2α must also be considered to avoid incomplete or potentially misleading interpretations. Clinically, this is particularly relevant as HIF-2α-targeting therapies such as Belzutifan become increasingly integrated into oncology practice. Defining how cytoplasmic HIF-2α contributes to tumor biology will be essential for identifying patients most likely to benefit from such treatments, while minimizing overtreatment and unnecessary side effects.

Although our conclusions are drawn from studies on tumor cells, the implications extend far beyond cancer. HIF-2α plays fundamental roles in numerous physiological contexts, including metabolic adaptation, oxygen sensing, tissue homeostasis, and embryonic development. Our work reveals that the biochemical regulation and functional repertoire of HIF-2α are far more complex than previously appreciated. A deeper understanding of these mechanisms will be critical for advancing our knowledge of early life development, organismal homeostasis, and disease.

## Materials and methods

### Cell culture, vector cloning, luciferase assays

SK-N-BE(2)c, SH-SY5Y, SH-EP, SK-N-SH, SK-N-AS, and 786-O cells were obtained from ATCC; SK-N-BE(2) from DSMZ. Cells were propagated in DMEM media, supplemented with 10% FBS (Thermo Fisher) and regularly tested for mycoplasma. Vectors used are listed in Supplementary Table S2. HIF-2α 2M P405A/P531A with N-terminal HA tag was a gift from William Kaelin (*41*) (Addgene plasmid #18956). pcDNA3.1-HA was a gift from Oskar Laur (Addgene plasmid #128034). Site-directed mutagenesis was used to create HIF-2α 3M variant P405A/P531A/N847A, and WT HIF-2α, and PCR mutagenesis was used to create HIF-2α deletion library. For luciferase assays, cells were co-transfected with HIF-2α variant, HRE-luciferase (*42*), and CMV-Renilla (Promega) vectors at a 1:1:0.08 ratio using Lipofectamine 3000 (Thermo Fisher). Luciferase signal was quantified 20-24 hours later using the Dual-Luciferase Reporter Assay (Promega). Signals were normalized to Renilla, with three technical replicates in each experiment. Cells from a fourth technical replicate in each experiment were extracted in 2x Laemmli buffer and immunoblotted to determine expression of the variants in each experiment. HRE-luciferase vector was a gift from Navdeep Chandel (Addgene #26731).

### Immunocytochemistry

Cells were transfected with HIF-2α variants using Lipofectamine 3000 (Thermo Fisher) for 24 hours, fixed with 4% PFA, permeabilized with 0.1% Triton-X100 and incubated with HA antibody (#901502, BioLegend) overnight. All antibodies used are listed in Supplementary Table S3. Cells were incubated with Alexa Fluor 568 secondary antibody (A-11004, Thermo Fisher), Phalloidin (65906, Sigma-Aldrich) and DAPI (D3571, Thermo Fisher) for 1 hour in room temperature. Serial Z-stacks were taken using Carl Zeiss LSM 780 laser scanning microscope with acquisition using ZEN Black 2012 software, using a 63X water immersion objective.

SK-N-BE(2)c cells were seeded onto glass coverslips at a density of 30,000 cells per well and cultured overnight at 37 °C. Cells were fixed the following day with 4% PFA, permeabilized with 0.3% Triton X-100, and blocked using serum-free protein blocking buffer (X090930-2, Agilent). Samples were then incubated overnight at 4°C with a primary antibody against TOMM20 (sc-17764, Santa Cruz Biotechnology), RPS6 (2317s, Cell Signalling Tech.), P4HB (ab2792, Abcam), or GORASP2 (66627-1, Proteintech). The next day, cells were incubated with alpaca anti-mouse Alexa Fluor 647 secondary antibody (615-605-214, Jackson ImmunoResearch) for 1 hour at room temperature, followed by overnight incubation at 4°C with a primary antibody against HIF-2α (A700-300, Thermo Fisher Scientific). On the third day, cells were incubated for 1 hour at room temperature with alpaca anti-rabbit Alexa Fluor 488 secondary antibody (611-545-215, Jackson ImmunoResearch) and DAPI nuclear stain (D3571, Thermo Fisher Scientific). Serial z-stack images were acquired using a Leica Stellaris 5 laser scanning confocal microscope with LAS X software and a 63× water-immersion objective. Representative images are shown as single focal planes, with brightness and contrast adjusted for visualization. Colocalization between HIF-2α and organelle marker signals was quantified using an in-house CellProfiler-based analysis pipeline, available upon request.

### Fractionation

For fractionation, cells were seeded and incubated in 21% or 1% O2 for 24 hours. Cells were lysed and fractionated using NE-PER Nuclear and Cytoplasmic Extraction Reagent (Thermo Fisher) according to the manufacturer’s protocol. Protein concentrations were measured using Bradford assay and equal amounts of cytoplasmic and nuclear fractions were loaded for Western blot. SDHA and Lamin B1 serve as cytoplasmic and nuclear markers, respectively.

### In vivo experiments

Female athymic mice (NMRI-Nu/Nu strain; Taconic) were housed in a controlled environment. 0.375*10^6^ SK-N-BE(2) or 0.75*10^6^ SK-N-AS cells were subcutaneously injected with a 100μl 2:1 mixture of cell culture media and growth factor-reduced Matrigel (BD, 354230) to the right flank of the mice. Mice were euthanized when the tumors reached a diameter of >1800 mm^3^ or at latest 230 days after the start of the experiment. All *in vivo* procedures followed the guidelines set by the Malmö-Lund Ethics Committee for the use of laboratory animals and were conducted in accordance with the European Union directive on the subject of animal rights (ethical permit no. 18743/19).

### Immunoprecipitation and sample processing for LC-MS/MS

For immunoprecipitation in proteomic experiments, four 15 cm dishes were seeded per IP with SK-N-BE(2) cells at a density of 8 x 10^6^ cells/dish in 24 ml of complete media. Next day, the cells were transfected with 30 μg of plasmids using Lipofectamine 3000 (Thermo Fisher) according to the manufacturer’s protocol. For hypoxia experiments, cells were cultured in a humidified chamber set to 1% O2 for 5 hours post-transfection and harvested 30 hours later. For normoxic (21% O2) experiments, cells were harvested approximately 30 hours after transfection. 786-O cells were seeded at a density of 3.6 x 10^6^ cells/dish and transfected the next day by 786-O Cell Avalanche Transfection reagent (EZ Biosystems); 9.25 μg plasmid per plate and 14.8 μl of the reagent. Medium was replaced 5 hours after transfection. Cells were washed with ice cold PBS (Gibco) and lysed for 30 minutes on ice with IP lysis buffer containing 50 mM Tris-Cl pH 7.4, 150 mM NaCl, 0.5% (v/v) NP-40, 0.25% (w/v) sodium deoxycholate, 2 mM EDTA, Complete Mini protease inhibitor cocktail tablets (Roche) and Pierce Phosphatase Inhibitor Mini Tablets (Thermo Fisher). Lysates were cleared by centrifugation (13000g, 4°C, 20 minutes) and FLAG (F1804, Sigma-Aldrich) or HA (#901502, BioLegend) antibodies were added at concentration 1μg/ 1ml lysate. Lysates were rotated at 4°C for 1 hour and immunocomplexes were then bound to Pierce ChIP-grade Protein A/G Magnetic Beads (160 μl/ 10ml of lysate) overnight. Lysates were separated into four low protein binding tubes (Thermo Fisher) that were used in all subsequent steps. Collected beads in each tube were washed 5x with 1ml of IP lysis buffer without inhibitors, and immunocomplexes were eluted using 0.1M glycine pH 2.3 for 5 minutes at RT and neutralized by equal amount of 0.2M ammonium bicarbonate pH 8. Protein concentration was measured by Bradford assay and 30 μg of eluates were reduced by adding 10 mM DTT (30 minutes, 56°C, with shaking), and alkylated by 20 mM iodacetamide (30 minutes, RT, in dark). Immunocomplexes were precipitated by adding 99.7% ethanol to a final concentration of 90% overnight at -20°C. Immunocomplexes were then centrifuged (14000g, 4°C, 20 minutes) and resolved in 100 μl of 0.1M ammonium bicarbonate pH 8. Trypsin (Sequencing Grade Modified Trypsin, Part No. V511A, Promega) was added in ratio 1:50 to protein concentration (0.6 μg trypsin / 30 μg sample) and samples were incubated overnight at 37°C. Next, 10 μl of trifluoroacetic acid (TFA) was added and samples were dried by centrifugal evaporator and stored at -80°C before MS analysis. Before MS the samples were resuspended in 0.1% TFA, 2% ACN. Peptide concentration was measured by NanoDrop (DeNovix DS-11, DeNovix Inc, Wilmington, USA) and 0.5 µg was injected to LC-MS/MS.

### LC-MS/MS Analysis

Samples were analyzed using two mass spectrometry platforms: the Exploris 480 (Thermo Fisher) coupled with a Vanquish Neo UHPLC system and the Eclipse Tribrid (Thermo Fisher) coupled with an Ultimate 3000 RSLC nano system. In both setups, a two-column configuration was used, where peptides were first loaded onto an Acclaim PepMap 100 C18 precolumn (75μm x 2 cm, Thermo Fisher, Waltham, MA) and subsequently separated on an EASY-Spray column. The columns were maintained at 45 °C, with a flow rate of 300 nL/min. Solvent A (0.1% FA in water) and solvent B (0.1% FA in 80% ACN) were used to create a nonlinear gradient to elute the peptides. For the Exploris system, separation occurred on a 25 cm EASY-Spray column (75 μm, C18, 2 μm, 100 Å, ES902) using a 60-minute nonlinear gradient: 5–25% solvent B over 50 minutes, increased to 32% over 6 minutes, and then to 45% over 4 minutes. For the Eclipse system, separation used a 50 cm EASY-Spray column (75 μm, C18, 2 μm, 100 Å, nanoViper). The gradient of solvent B was maintained at 3% for 4 min, then increased to 25% over 70 minutes, increased to 38% over 20 minutes, and then to 95% over 1 minute. Finally, the gradient was held for 5 minutes for column washing. Both systems used data-dependent acquisition (DDA) in positive ion mode. The Exploris operated with an MS1 resolution of 120,000 (m/z 200), a normalized AGC target of 300%, and a maximum injection time of 45 ms, with a mass range of 375–1500 m/z. The Eclipse employed an MS1 resolution of 120,000 in the normal mass range, with a standard AGC target and automatic maximum injection time, spanning 350–1400 m/z. Precursor ions were isolated with a window of 1.3 m/z for Exploris and 1.6 m/z for Eclipse, fragmented by HCD at a normalized collision energy of 30, and detected at 15,000 resolution in the Orbitrap. Dynamic exclusion settings were 30 seconds for Exploris.

### MS Data analysis

DDA files were processed using Proteome Discoverer™ 2.5 (Thermo Scientific), and peptide identification was performed with SEQUEST HT against the UniProtKB human reference proteome (UP000005640_9606). Searches included carbamidomethylation of cysteine as a static modification and N-terminal acetylation and methionine oxidation as dynamic modifications. Precursor and fragment mass tolerances were set to 10 ppm and 0.02 Da, respectively, and up to two missed cleavages were allowed. Peptide-spectrum matches were validated using Percolator with a maximum q-value threshold of 0.05.

Reproducibility across experiments was evaluated by correlating peptide counts for individual proteins in HIF-2α immunoprecipitations. The resulting R² matrix (Supplementary Fig. S4C) indicated high intra-group concordance across SK-N-BE(2) normoxic, SK-N-BE(2) hypoxic, and 786-O replicates. To minimize nonspecific binders, proteins were provisionally classified as candidate interactors if their peptide count ratio (IP/Ctrl) exceeded 2 or if no peptides were detected in the corresponding control sample. In total, 29 immunoprecipitation experiments were included: 11 normoxic SK-N-BE(2) replicates (including four from the SK-N-BE(2)c subclone), nine hypoxic (1% O₂) SK-N-BE(2) replicates, and nine normoxic 786-O replicates. These yielded 2,663, 2,946, and 1,238 non-redundant preliminary candidate proteins, respectively.

To derive high-confidence interactomes, we applied replicate-detection thresholds tailored to dataset quality, as determined by correlation analysis (Supplementary Fig. S4C). Proteins were retained if classified as candidate interactor in at least 5 of 11 normoxic SK-N-BE(2) experiments, 7 of 9 hypoxic SK-N-BE(2) experiments, or 4 of 9 786-O experiments. This filtering strategy resulted in 135 interactors in 786-O cells, and 168 in normoxic and 241 in hypoxic SK-N-BE(2) cells. Complete interactor lists for all conditions are provided in Supplementary Table S1. For Fig. 6E, quantitative proteomics data were imported into R and filtered to retain proteins shared across all three experimental conditions. Log2 fold changes between HIF overexpression and empty vector controls were calculated using a pseudocount of 1 to avoid division by zero. Heatmaps were generated using the pheatmap package with hierarchical clustering, and samples were grouped by experimental condition.

### Gene ontology and STRING analysis

List of interactors from MS results were loaded into R v4.2.2. Gene ontology enrichment analysis was performed for Biological Processes (BP), Molecular Function (MF) and Cellular Compartment (CC) using clusterProfiler v4.14.4, with Org.Hs.eg.db v3.20.0 as the annotation database. Significantly enriched GO terms were identified with a Benjamini-Hochberg correction (p<0.05) and plotted using clusterProfiler and ggplot2 v3.5.1.

Protein-protein interaction networks were generated using the STRING database. Protein lists derived from mass spectrometry experiments and classified as normoxic, cell type-independent or oxygen- and cell type-independent were uploaded to STRING to evaluate known and predicted functional associations among detected proteins. Networks were generated using a minimum required interaction score of 0.700, corresponding to high-confidence interactions.

For protein-interaction network analyses in Supplementary Figs. S5-S7, the three HIF-2α interactor sets were queried in the stringApp for Cytoscape using the following settings: species = Homo sapiens, physical interaction network, confidence score = 0.8, and no additional interactors. Nodes were grouped based on physical interaction, shared biological processes, or subcellular localization.

### RNA sequencing

RNA was extracted using the Qiagen RNeasy Mini Kit according to the manufacturer’s instructions. RNA quality and concentration were assessed using a TapeStation 4200 (Agilent) and Qubit Flex fluorometer (Thermo Fisher Scientific). Libraries were prepared from ∼250 ng total RNA using the Illumina Stranded mRNA Prep kit with 13 PCR amplification cycles and unique dual indexing. Library quality and concentration were evaluated using TapeStation and Qubit. Sequencing was performed on an Illumina NovaSeq 6000 platform to generate paired-end 150 bp reads with 1% PhiX spike-in. Raw reads were trimmed using fastp, and ribosomal RNA reads were removed. Reads were aligned to the human GRCh38 reference genome using STAR, and transcript quantification was performed with Salmon. Gene-level counts were generated using featureCounts based on GENCODE v45 annotation. Quality control was performed using standard tools implemented in the nf-core/rnaseq pipeline. Downstream analyses were conducted using variance-stabilized counts from DESeq2. Differential expression analysis was performed using DESeq2. Due to pronounced batch effects identified by principal component analysis and hierarchical clustering, samples from replicate 2 were excluded from downstream analyses. Differentially expressed genes were defined as those with an adjusted p-value < 0.05 and absolute log2 fold change > 1.

### Statistical analyses and quantifications

Correlations among HRE transactivation, protein expression, and nuclear and cytoplasmic localization (Fig. 3N, using data from Fig. 3M) were assessed by nonparametric Spearman correlation analysis in GraphPad Prism v10.5.0. Spearman correlation coefficients (*r*) were calculated, and statistical significance was determined using two-tailed *P* values with 95% confidence intervals. Western blots were quantified using ImageJ by averaging three different exposures per independent experiment, the HIF-α signals were normalized to loading control. For calculating the expression levels of the HIF-2α variants in Fig. 3I-L, these densitometry values were used, from which the empty vector control values were subtracted, and relative values where HIF-2α was set to one were calculated. The values for normalization of the luciferase assays from the Western blots were calculated equally. The luciferase values were first normalized to Renilla and then to HIF-2α (HIF-2α = 1). These values were normalized to HIF-2α relative expression data (where HIF-2α = 1).

## Data Availability

The mass spectrometry proteomics data have been deposited to the ProteomeXchange Consortium via the PRIDE partner repository with the dataset identifier PXD059290. The RNA sequencing data have been deposited in the GEO database and are accessible under accession number GSE329133.

## Acknowledgements

The authors would like to acknowledge Clinical Genomics Lund, SciLifeLab and Center for Translational Genomics (CTG), Lund University, for providing expertise and service with sequencing and analysis. The authors would further like to acknowledge Charlotte Welinder for her valuable contributions to experimental proteomics design. This study was funded by the Swedish Cancer Society, the Swedish Childhood Cancer Fund, the Crafoord Foundation, Mrs. Berta von Kamprad Foundation, the Ollie and Elof Ericsson Foundation, the Magnus Bergvall Foundation, the Hans von Kantzow Foundation, and the Royal Physiographic Society of Lund.

## Author contributions

Conceptualization: TG, SM

Methodology: TG, SK, AZ, SM

Software: AZ, SÖ

Validation: TG, SK

Formal analysis: TG, AZ, SÖ

Investigation: TG, SK, AZ, EF, SÖ, CW, JH

Data curation: TG, SM

Writing - original draft: TG, JR, SM

Writing - review & editing: TG, SK, AZ, SÖ, EF, CW, EH, JH, JR, SM

Visualization: TG, SK, AZ, SM

Supervision: SM Funding acquisition: SM

Project administration: SM

