## Supplementary Material for "Structural survey of HIF-2α reveals regulation of its subcellular localization and protein interactome"

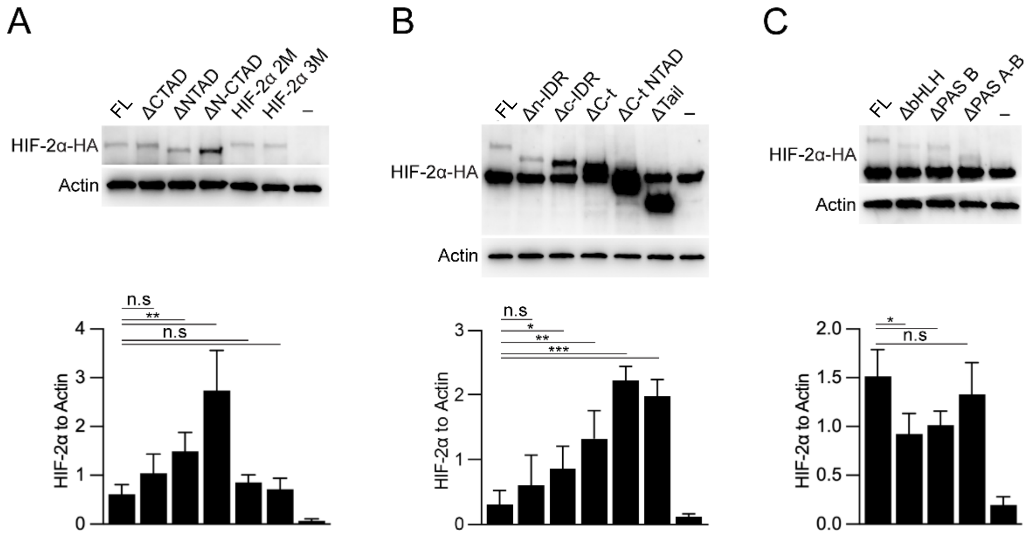


**Figure S1**. *Expression of HIF-2α variants in hypoxia and their subcellular localization in normoxia.* **(A–C)** Expression of HIF-2α variants in neuroblastoma SK-N-BE(2)c cells cultured at 1% $O_2$. Western blot signal was quantified, normalized to actin values, and graphed from *n*=4 independent experiments (Student's t-test; *n.s*; non-significant, * *p* <0.05, ** *p* <0.01, *** *p* <0.001). FL; full-length HIF-2α variant.

Supplementary Figure S2

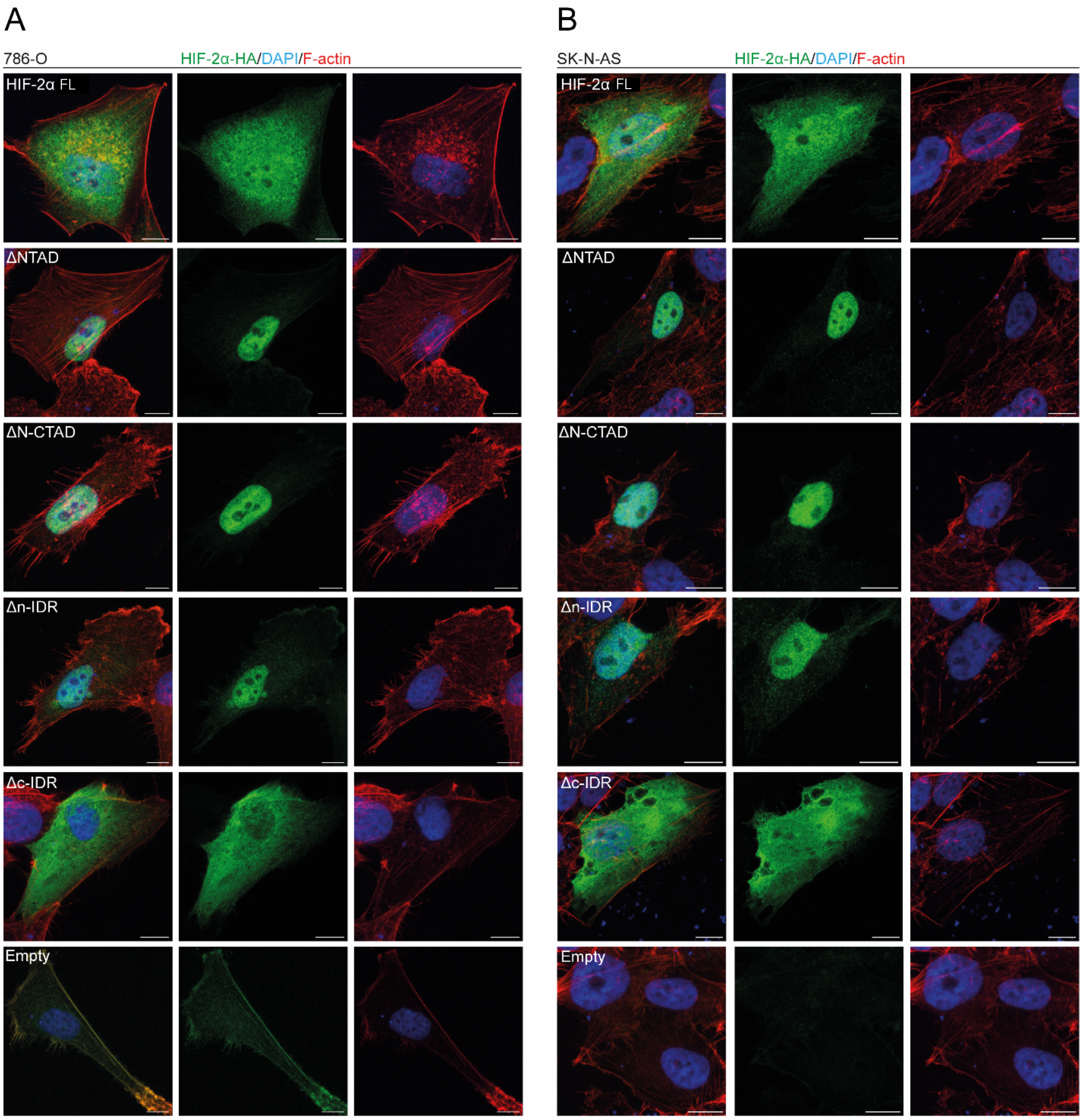


**Figure S2.** *HIF-2α variants display similar localization patterns in neuroblastoma and renal cell carcinoma cells.* **(A–B)** The HA-tagged HIF-2α variants were expressed in renal cell carcinoma 786-O (**A**) and neuroblastoma SK-N-AS (**B**) cells, and immunolabeled for HA epitope. F-actin was counterstained with Phalloidin (red), and DAPI (blue) was used to visualize nuclei. *n*=3 independent experiments; scale bars, 10 µm. FL; full-length HIF-2α variant.

Supplementary Figure S3

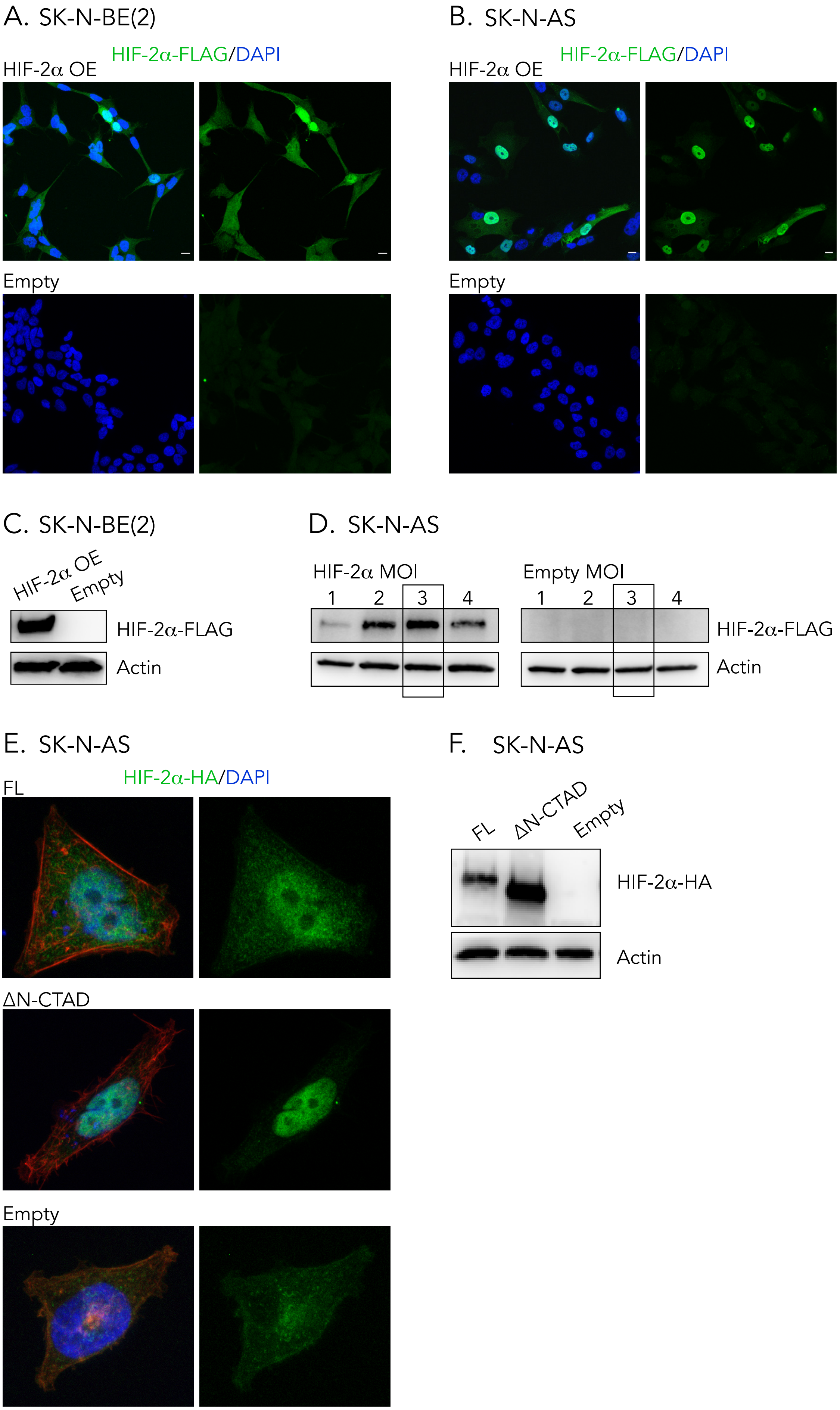

**Figure S3.** *Validation of HIF-2α expression in cells used for in vivo experiments.* **(A–B)** Immunocytochemical staining for FLAG (green) in neuroblastoma SK-N-BE(2) (**A**) and SK-N-AS (**B**) cells transduced with empty vector or a FLAG-tagged HIF-2α overexpression construct. Nuclei were visualized with DAPI (blue). **(C)** Western blot analysis of FLAG expression in SK-N-BE(2) cells transduced with empty vector or FLAG-tagged HIF-2α. **(D)** Western blot analysis of FLAG expression in SK-N-AS cells transduced with empty vector or FLAG-tagged HIF-2α. Cell populations transduced at different multiplicities of infection (MOI) are shown; the population used for *in vivo* experiments is indicated by a boxed outline. **(E)** Immunocytochemical staining for HA (green) in SK-N-AS cells transfected with empty vector or HA-tagged HIF-2α full-length (FL) or ΔN-CTAD constructs. F-actin was counterstained with phalloidin (red), and nuclei were visualized with DAPI (blue). **(F)** Western blot validation of HIF-2α expression in the same cell populations shown in (**E**). Actin was used as a loading control in all immunoblot analyses.

Supplementary Figure S4

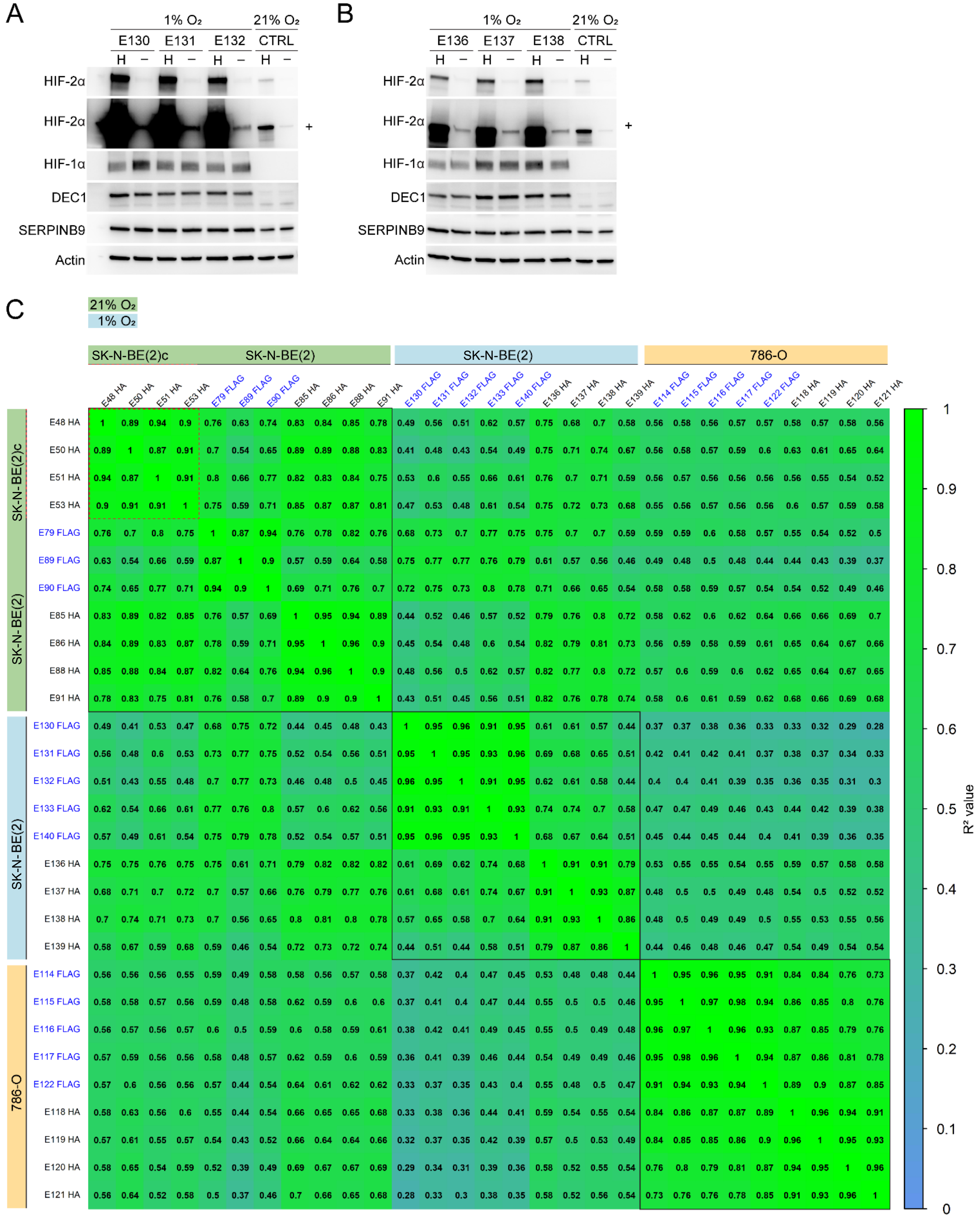

**Figure S4.** *Validation of the oxygen response in hypoxic IP-MS experiments.* **(A–B)** Expression levels of HIF-1α, HIF-2α and known hypoxia-driven proteins DEC1 and SERPINB9 in neuroblastoma SK-N-BE(2) cells grown at 1% $O_2$, as assessed by Western blot. Actin was used as loading control. E130-132 indicate *n*=3 independent FLAG IP experiments (**A**), E136-138 indicate *n*=3 independent HA IP experiments (**B**). H = HIF-2α overexpression lysate (IP); – = Empty lysate (control); CTRL – control cells transfected and grown in 21% $O_2$; + indicates stronger exposure of the upper HIF-2α blot. **(C)** Correlation plot of all IP-MS experiments. Correlations ($R^2$) of detected peptide counts per protein in HIF-2α immunoprecipitation samples are shown. Black squares highlight experiments from each group; red dotted line indicate experiments in SK-N-BE(2)c cells.

Supplementary Figure S5

SK-N-BE(2); 1% $O_2$

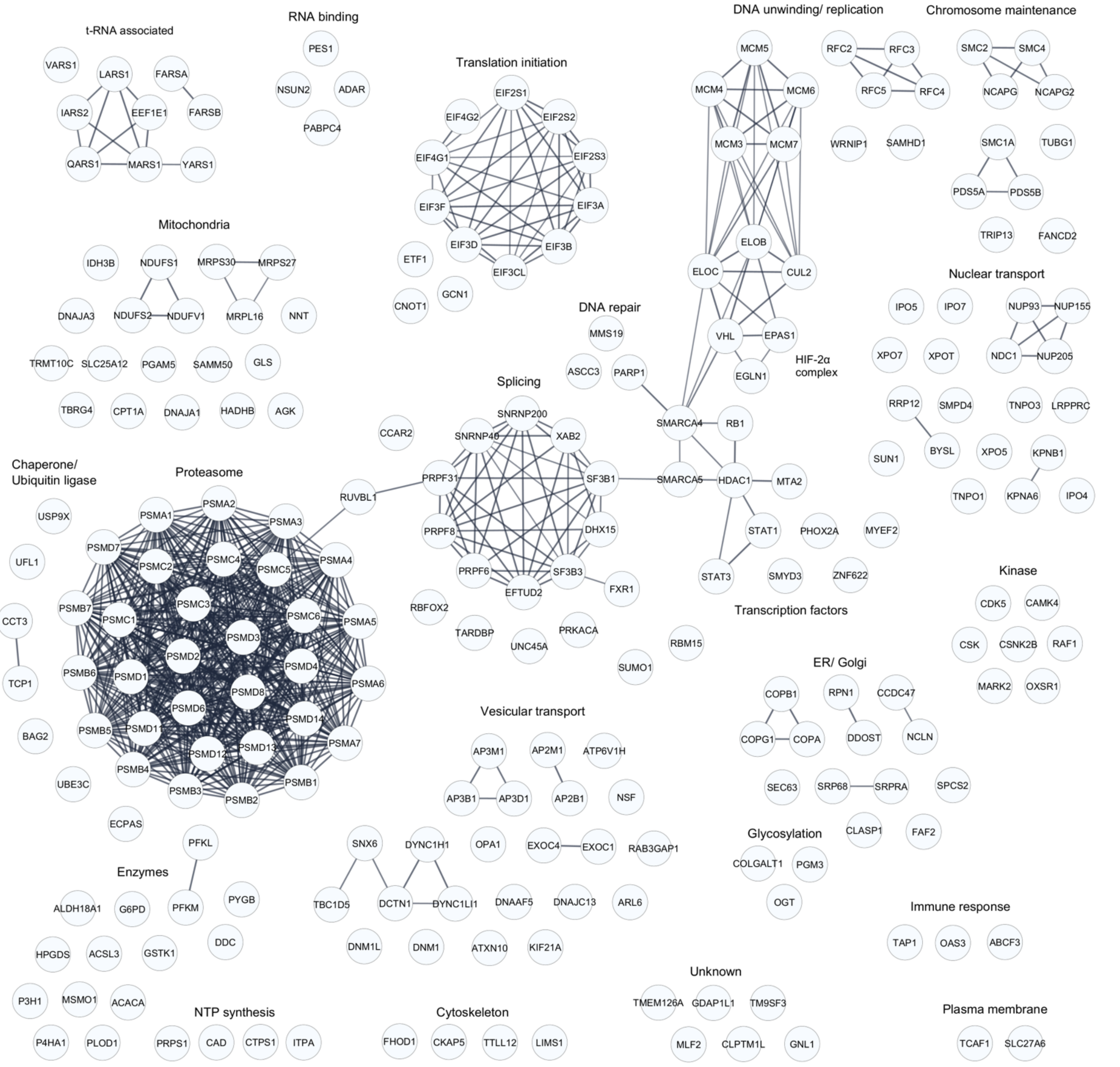


**Figure S5.** *STRING analysis of HIF-2α interactors in neuroblastoma SK-N-BE(2) cells cultured at 1% $O_2$.* Network of known physical protein interactions within identified proteins. Only high-confidence interactions are shown as connections (STRING cutoff = 0.8).

Supplementary Figure S6

SK-N-BE(2); 21% $O_2$

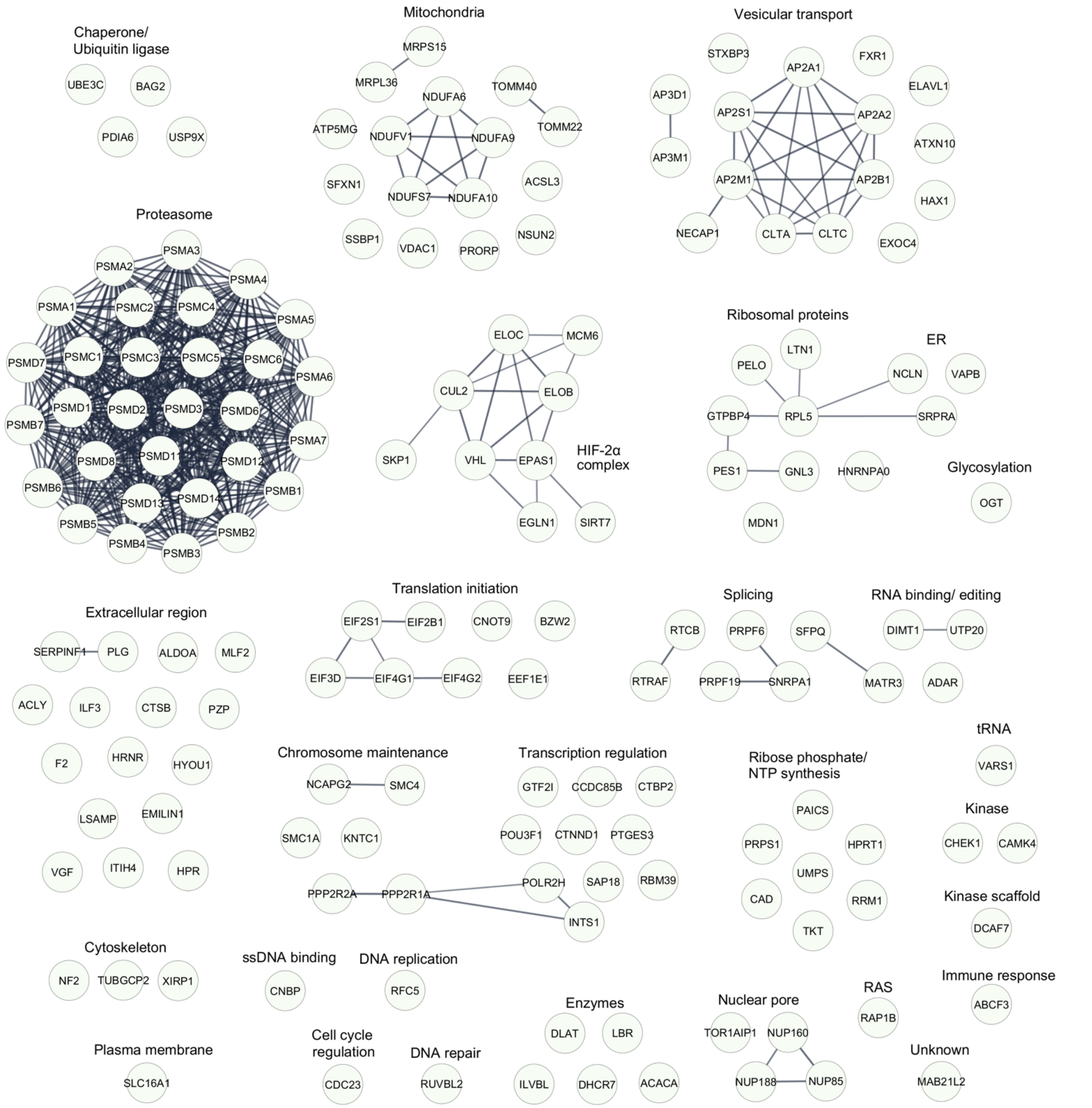


**Figure S6.** *STRING analysis of HIF-2α interactors in SK-N-BE(2) cells cultured at 21% $O_2$.* Network of known physical protein interactions within the identified proteins. Only high-confidence interactions are shown as connections (STRING cutoff = 0.8).

Supplementary Figure S7

786-O; 21% $O_2$

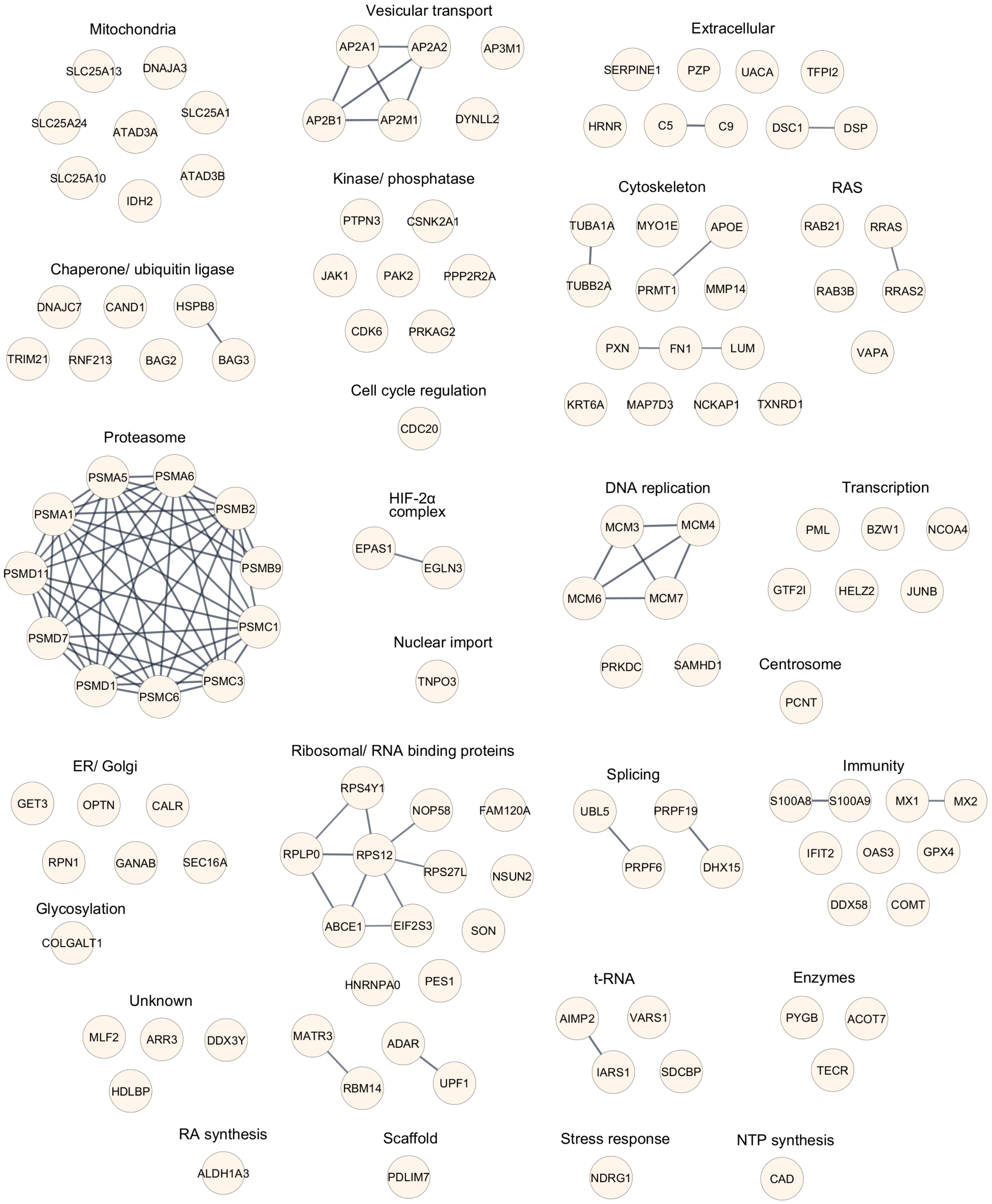


**Figure S7.** *STRING analysis of HIF-2α interactors in 786-O cells cultured in 21% $O_2$.* Network of known physical protein interactions within identified proteins. Only high-confidence interactions are shown as connections (STRING cutoff = 0.8).

Supplementary Table S1.

Complete data from IP-MS. Comprehensive lists of HIF-2α interactors identified across three experimental conditions: SK-N-BE(2) neuroblastoma cells cultured at 21% or 1% $O_2$, and 786-O ccRCC cells cultured at 21% $O_2$. Details on HIF-2α coverage, mass spectrometers used, and experimental runs are provided. Interactors are ranked from most to least frequently detected across all experiments. "**Candidate sum in all exp**" indicates the total number of experiments in which each protein was classified as an interactor, and "**% of all exp**" shows the corresponding percentage. "**Candidate #**" denotes whether the protein was present (=1) or absent (=0) in each experiment; green and blue shading represent HA-tag and FLAG-tag immunoprecipitations, respectively. "**Literature**" indicates whether the interactor is previously reported in **BioGRID**; interactors from Daly *et al.* (2021) (*1*), not yet curated in BioGRID, were manually added. For SK-N-BE(2) 21% $O_2$, hypoxic interactors from Daly *et al.* were excluded; for SK-N-BE(2) 1% $O_2$, normoxic interactors were excluded; the 786-O dataset was cross-referenced against the full Daly *et al.* list. The bold horizontal line marks the threshold used to define candidate interactors. Additional columns summarize experiment-specific analyses. "**Candidate final if = 2**" merges the values from "Candidate ratio" and "Candidate specific H>0 and E=0"; a summed value >2 designates the protein as a candidate in that experiment. "**Candidate ratio**" =1 when the HIF-2α/Empty peptide ratio >2. "**Candidate specific H>0 and E=0**" returns 2 if both conditions are met and 1 if only one is met. "**#peptides_H**" and "**#peptides_E**" list the number of peptides detected in HIF-2α and Empty controls, respectively.

Separate Excel file.

Supplementary Table S2. Vectors used in this study.

| # | Plasmid | Tag | Gene | Sequence | Backbone | Origin |
|---|---|---|---|---|---|---|
| 1 | pcDNA3.1 HA-HIF-2α WT | HA (N-terminal) | human EPAS1/HIF-2α | 1-870 (WT) | pcDNA3.1 | Cloned from HA-HIF-2α 2M |
| 2 | pcDNA3.1 HA-ΔCTAD | HA (N-terminal) | human EPAS1/HIF-2α | 1-820 | pcDNA3.1 | Cloned from HA-HIF-2α WT |
| 3 | pcDNA3.1 HA-ΔNTAD | HA (N-terminal) | human EPAS1/HIF-2α | Δ496-542 | pcDNA3.1 | Cloned from HA-HIF-2α WT |
| 4 | pcDNA3.1 HA-ΔN-CTAD | HA (N-terminal) | human EPAS1/HIF-2α | Δ496-542, Δ820-870 | pcDNA3.1 | Cloned from HA-HIF-2α WT |
| 5 | pcDNA3.1 HA-Δn-IDR | HA (N-terminal) | human EPAS1/HIF-2α | Δ362-485 | pcDNA3.1 | Cloned from HA-HIF-2α WT |
| 6 | pcDNA3.1 HA-Δc-IDR | HA (N-terminal) | human EPAS1/HIF-2α | Δ578-819 | pcDNA3.1 | Cloned from HA-HIF-2α WT |
| 7 | pcDNA3.1 HA-ΔC-t | HA (N-terminal) | human EPAS1/HIF-2α | 1-577 | pcDNA3.1 | Cloned from HA-HIF-2α WT |
| 8 | pcDNA3.1 HA-ΔC-t NTAD | HA (N-terminal) | human EPAS1/HIF-2α | 1-485 | pcDNA3.1 | Cloned from HA-HIF-2α WT |
| 9 | pcDNA3.1 HA-ΔTail | HA (N-terminal) | human EPAS1/HIF-2α | 1-361 | pcDNA3.1 | Cloned from HA-HIF-2α WT |
| 10 | pcDNA3.1 HA-ΔbHLH | HA (N-terminal) | human EPAS1/HIF-2α | Δ13-67 | pcDNA3.1 | Cloned from HA-HIF-2α WT |
| 11 | pcDNA3.1 HA-ΔPAS B | HA (N-terminal) | human EPAS1/HIF-2α | Δ241-351 | pcDNA3.1 | Cloned from HA-HIF-2α WT |
| 12 | pcDNA3.1 HA-ΔPAS A-B | HA (N-terminal) | human EPAS1/HIF-2α | Δ84-351 | pcDNA3.1 | Cloned from HA-HIF-2α WT |
| 13 | pcDNA3.1 HA-HIF-2α 2M | HA (N-terminal) | human EPAS1/HIF-2α | P405A, P531A | pcDNA3.1 | Addgene #18956 |
| 14 | pcDNA3.1 HA-HIF-2α 3M | HA (N-terminal) | human EPAS1/HIF-2α | P405A, P531A, N847A | pcDNA3.1 | Cloned from HA-HIF-2α 2M |
| 15 | pcDNA3.1 HA-empty | HA (N-terminal) | Not applicable | Not applicable | pcDNA3.1 | Addgene #128034 |
| 16 | pGL2 HRE luciferase | Not applicable | Luciferase | Not applicable | pGL2 | Addgene #26731 |
| 17 | pRL CMV-Renilla luciferase | Not applicable | Renilla luciferase | Not applicable | pRL | Promega (cat. no. E6931) |

Supplementary Table S3. Antibodies used in this study

| Target | Application | Cat. No. | Vendor |
|---|---|---|---|
| HIF-2α | WB/ ICC | A700-003 | Thermo Fisher Scientific |
| HIF-1α | WB | 610959 | BD Biosciences |
| HA.11 | IP/ WB/ ICC | 901502 | BioLegend |
| FLAG | IP/ WB | F1804 | Sigma-Aldrich |
| SDHA | WB | ab14715 | Abcam |
| Lamin B1 | WB | 12586 | Cell Signaling Technology |
| Actin | WB | 0869100-CF | MP Bio |
| DEC1 | WB | NB100-1800 | Novus Bio |
| VEGF | WB | sc-7269 | Santa Cruz |
| SERPINB9 | WB | PA5-92943 | Thermo Fisher Scientific |
| Goat anti-Mouse IgG AF 568 | ICC | A-11004 | Thermo Fisher Scientific |
| RPS6 | ICC | 2317s | Cell Signalling Tech |
| P4HB (PDI) | ICC | ab2792 | Abcam |
| GORASP2 | ICC | 66627-1 | Proteintech |
| TOMM20 | ICC | sc-17764 | Santa Cruz |
| alpaca anti-mouse Alexa Fluor 647 | ICC | 615-605-214 | Jackson ImmunoResearch |
| alpaca anti-rabbit Alexa Fluor 488 | ICC | 611-545-215 | Jackson ImmunoResearch |
